# Effects of disease emergence on invasive grass impacts

**DOI:** 10.1101/2022.03.09.483680

**Authors:** Amy E. Kendig, Ashish Adhikari, Brett R. Lane, Christopher M. Wojan, Nicholas Kortessis, Margaret W. Simon, Michael Barfield, Philip F. Harmon, Robert D. Holt, Keith Clay, Erica M. Goss, S. Luke Flory

## Abstract

Invasive species impact ecosystems through their large abundances and strong per capita effects. Enemies can regulate abundances and per capita effects, but are notably absent for many new invaders. However, invaders acquire enemies over time and as they spread; processes hypothesized to mitigate negative invader impacts by reducing abundance or per capita effects. Alternatively, properties of invaders or acquired enemies, such as an enemy’s ability to attack multiple species, may hinder enemy mitigation of invader impacts. We used field experiments to evaluate disease mitigation of invader impacts using the invasive grass *Microstegium vimineum*, which hosts an emerging fungal disease, and a native grass competitor, *Elymus virginicus*. We manipulated competition through density gradients of each plant species, and we reduced ambient foliar diseases with fungicide and autoclaving. We then modeled long-term population dynamics with field-estimated parameters. In the field, disease did not reduce invader abundance or per capita effects. The invader amplified disease on itself and the competitor, and disease reduced invader and competitor fitness components (e.g., germination). The dynamical model predicted that disease impacts on the competitor are greater than on the invader, such that disease will reduce invader abundance by 18%, and competitor abundance by 88%, over time. Our study suggests that enemies acquired by invaders will not necessarily mitigate invader impacts if the invader amplifies the enemy and the enemy attacks and suppresses competitor species.

## Introduction

Non-native species are frequently introduced to new locations through human-assisted pathways (1). A subset of these species become invasive and cause negative ecological impacts (2, 3). Substantial ecological impacts occur when invaders are abundant and have strong per capita effects on other species (Fig. 1A) (4, 5). Theory predicts that natural enemies can mitigate invader impact (Fig. 1B–C). For example, the enemy release hypothesis posits that invasive species can establish and outcompete native species because they lack co-evolved specialist enemies, which would otherwise reduce their abundance or per capita effects (6). Biological control is based on the premise that introducing specialist enemies reduces invader impacts (7). Most recently, the pathogen accumulation and invasive decline hypothesis suggests that enemies acquired during invasion (i.e., accumulated enemies) also mitigate invader impacts (8). Invaders accumulate enemies the longer they are established and the farther they spread (9, 10). Yet, there are few tests of whether and how accumulated enemies mitigate invader impacts (11).

**Figure 1.**
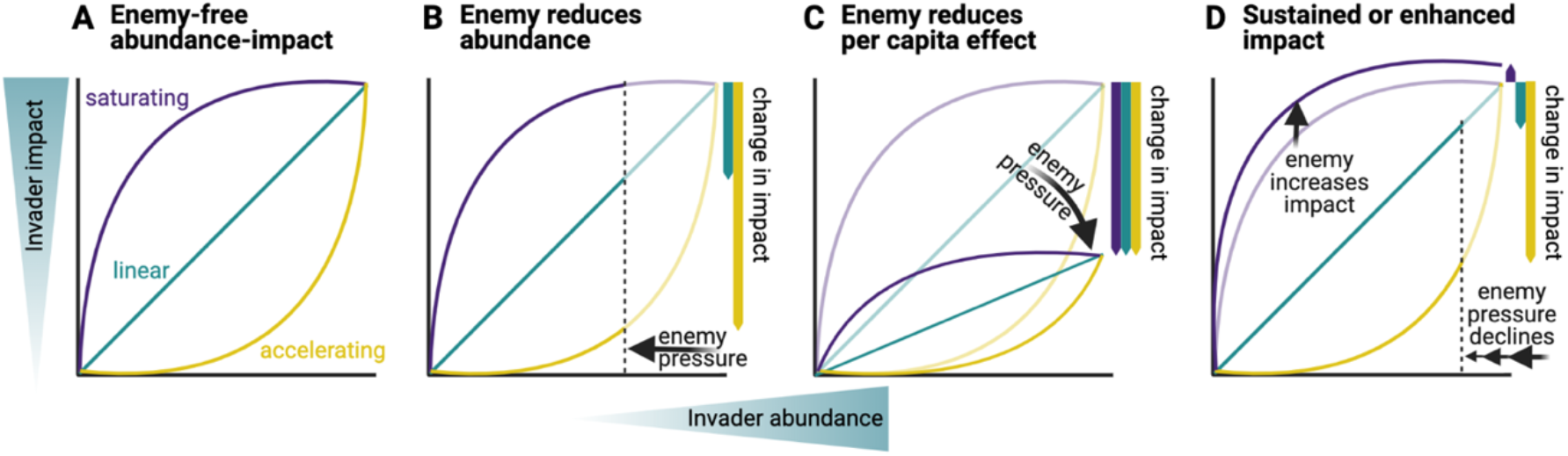
Potential enemy effects on invader abundance–impact relationships. (A) Larger invader abundance generally leads to greater impact, but the shape of the relationship may vary (shown here: linear (blue), saturating (purple), accelerating (yellow)). The per capita effect determines the change in impact with each unit increase in abundance. (B) Enemy pressure may reduce invader abundance, as represented by dark lines relative to light lines. Reduced abundance may reduce impact (linear or accelerating) or have no effect on impact (saturating). (C) Enemies may reduce per capita effects of invaders, as represented by dark lines relative to light lines. Reduced per capita effects may reduce impact (all three). (D) The predictions in B and C may not be realized because the enemy increases the impact (saturating) or because enemy pressure declines with invader abundance (linear or accelerating). Note: change in impact at maximum abundance relative to the enemy-free scenario in A is depicted on the right sides of panels B–D.

An important impact of species invasions is competitive suppression of native species (2, 3). Successful biological control suggests that competitors can recover with enemy-driven declines in invader abundance (12, 13). However, a saturating abundance–impact curve (Fig. 1A) may lead to sustained impact despite reductions in invader abundance (Fig. 1B) (14). Further, invasive species may tolerate enemy attacks, either at the scale of an individual (e.g., a weak effect of enemy damage on fitness) (15) or population (e.g., compensatory population growth following attacks) (16), limiting enemy effects on invader abundance altogether.

In contrast to biological control agents and specialist enemies featured in the enemy release hypothesis, generalist enemies acquired by invasive species may attack co-occurring species, potentially leading to apparent competition (17). Generalist enemies that spill over from co-occurring species may effectively reduce invader abundance and impact (Fig. 1B) (18, 19). However, spillback of enemies from the invader to other species may lead to sustained or increased invader impact despite reductions in invader abundance (Fig. 1D) (20, 21). Additionally, enemies that depend on invader abundance for persistence or dispersal (e.g., density-dependent disease transmission) may exert less pressure as an invader declines (Fig. 1D). Therefore, characterizing how much enemies depend on and affect invaders and competitors is crucial to understanding enemy effects on the invader abundance–impact relationship.

In addition to reducing invader abundance, enemies may also mediate invader impacts by altering invader per capita effects on competitors (Fig. 1C). To evaluate the net per capita effects of invaders, all types of competition, including apparent, resource, and interference, must be considered together (22). Theory predicts that if the net intraspecific effects of competing species exceed their net interspecific effects (i.e., they limit themselves more than others), they can potentially coexist. Otherwise, a species with greater interspecific than intraspecific effects may exclude its competitor, or priority effects may occur (22, 23). Intra- and interspecific effects of the invader and competitors provide insights into invader impacts because they can be linked to the abundance–impact relationship; intraspecific effects influence invader abundance and interspecific effects are synonymous to per capita effects.

We hypothesized that enemy accumulation mitigates invader impacts through reduced abundance or per capita effects, unless properties of the invader or enemies (e.g., saturating abundance–impact curve, density dependence, spillback) hinder this process. To test this hypothesis, we conducted field experiments with the invasive annual grass *Microstegium vimineum* (stiltgrass). The recent emergence of foliar fungal pathogens on *M. vimineum* in its introduced range (eastern and midwestern U.S.) may alter impacts of this invader on native plants (24, 25). “Emergence” here refers to pathogens that have been recently recognized on a host and have increased in incidence over time; in this case, fungi in the genus *Bipolaris* that were first detected on *M. vimineum* in the past 20 years (25). In the absence of *Bipolaris, M. vimineum* suppresses other plant species through resource competition and litter build-up (26, 27). The competitor in our study is a native perennial bunchgrass (*Elymus virginicus*) that commonly co-occurs with *M. vimineum* (28) and is susceptible to *Bipolaris* infection (29, 30). As a perennial, *E. virginicus* occurs as first-year seedlings and older adults, which may differ in their interactions with *M. vimineum*.

Here, we present the results from field experiments in forest in Indiana, USA. We manipulated invader and competitor abundances and disease (i.e., we reduced disease by spraying plants with a fungicide that suppresses *Bipolaris*, and by autoclaving litter). First, we measured short-term (within growing season) effects of disease on invader abundance and per capita effects. Then, we assessed how much disease depends on and affects fitness components of the invader and competitor. We conducted two repetitions of the field experiments over two years and results presented here are from second repetitions, unless otherwise noted (Methods S1). Finally, we modelled long-term population dynamics to evaluate how disease-induced changes in intraspecific and interspecific effects influence competitor abundance (i.e., invader impacts).

## Results

We found limited (non-statistically significant) evidence that disease reduced invader abundance over one growing season. Reducing disease increased invader biomass by 36% [-51%, 186%] (model-estimated median, 95% credible intervals in brackets, applies throughout) at high invader density (Fig. 2A, Table S1). We did not find evidence that reducing disease affected invader seed production (Fig. 2A, Table S2).

**Figure 2.**
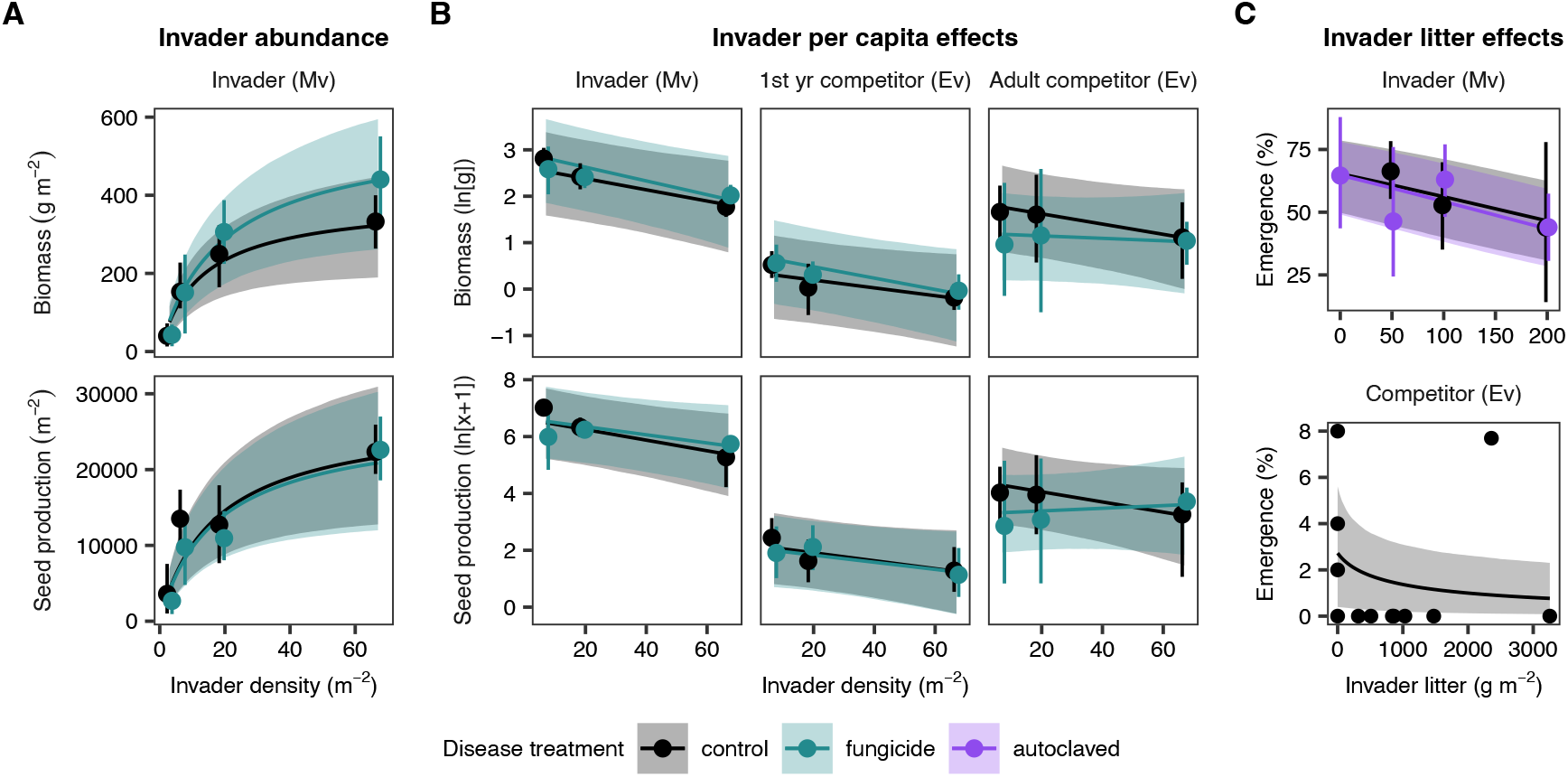
The effect of foliar fungal disease suppression on invader (*M. vimineum*, abbreviated as Mv) abundance, per capita effects on the invader and competitor (*E. virginicus*, abbreviated as Ev), and litter effects on both species. (A) Invader abundance was estimated with plot-scale biomass and seed production across a gradient of invader density. (B) Per capita effects were measured as change in biomass and seed production (both natural log-transformed) across the invader density gradient. (C) Litter effects were measured as change in percentage seeds emerged (emergence) across litter abundance gradients. Invader and competitor panels in C differ in invader litter gradients and disease treatments because they are two different experiments (see Methods for details). Points and error bars represent mean ± 95% bootstrapped confidence intervals of raw data, except the competitor panel of C where points represent raw data. Lines and shading represent model-estimated mean ± 95% credible intervals.

There was limited evidence that disease increased per capita effects of the invader. Reducing disease reduced the per capita effect of the invader on adult competitor growth by 79% [-456%, 338%] and seed production by 121% [-626%, 334%] (Fig. 2B, Tables S3–S6). Reducing disease did not alter per capita effects of the invader on its own or first-year competitor biomass or seed production (Fig. 2B, Tables S3–S6). Similarly, there was limited evidence that disease increased per capita effects of the invader in the first repetition of the experiment (Fig. S1, Table S7). There was strong evidence that invader litter reduced invader and competitor emergence (Fig. 2C, Tables S8–S9). Autoclaving litter did not alter the effect of litter on invader emergence (Table S8).

There was strong evidence that disease transmission depended on both the invader and competitor. Earlier in the growing season (8–12 weeks post planting), competitor plants transmitted disease to invaders in control plots and to first-year competitors in both treatments (Fig. 3, Table S10). Additionally, invader plants surrounding plots transmitted disease to invaders in fungicide-treated plots and first-year competitors in control plots (Fig. 3). Later in the growing season (16 weeks post planting), invader plants surrounding plots transmitted disease to invaders in both treatments, and invader plants within plots transmitted disease to competitors in control plots (Fig. 3). In the first repetition of the experiment, we only observed intraspecific transmission (Fig. S2, Table S11). Disease severity was somewhat surprisingly not related to environmental variation among plots (including dew intensity, soil moisture, and canopy cover), except that disease severity of first-year competitors declined with more dew intensity and soil moisture (Tables S12–S13). We identified *Bipolaris* fungi on leaves of both species (Table S14).

**Figure 3.**
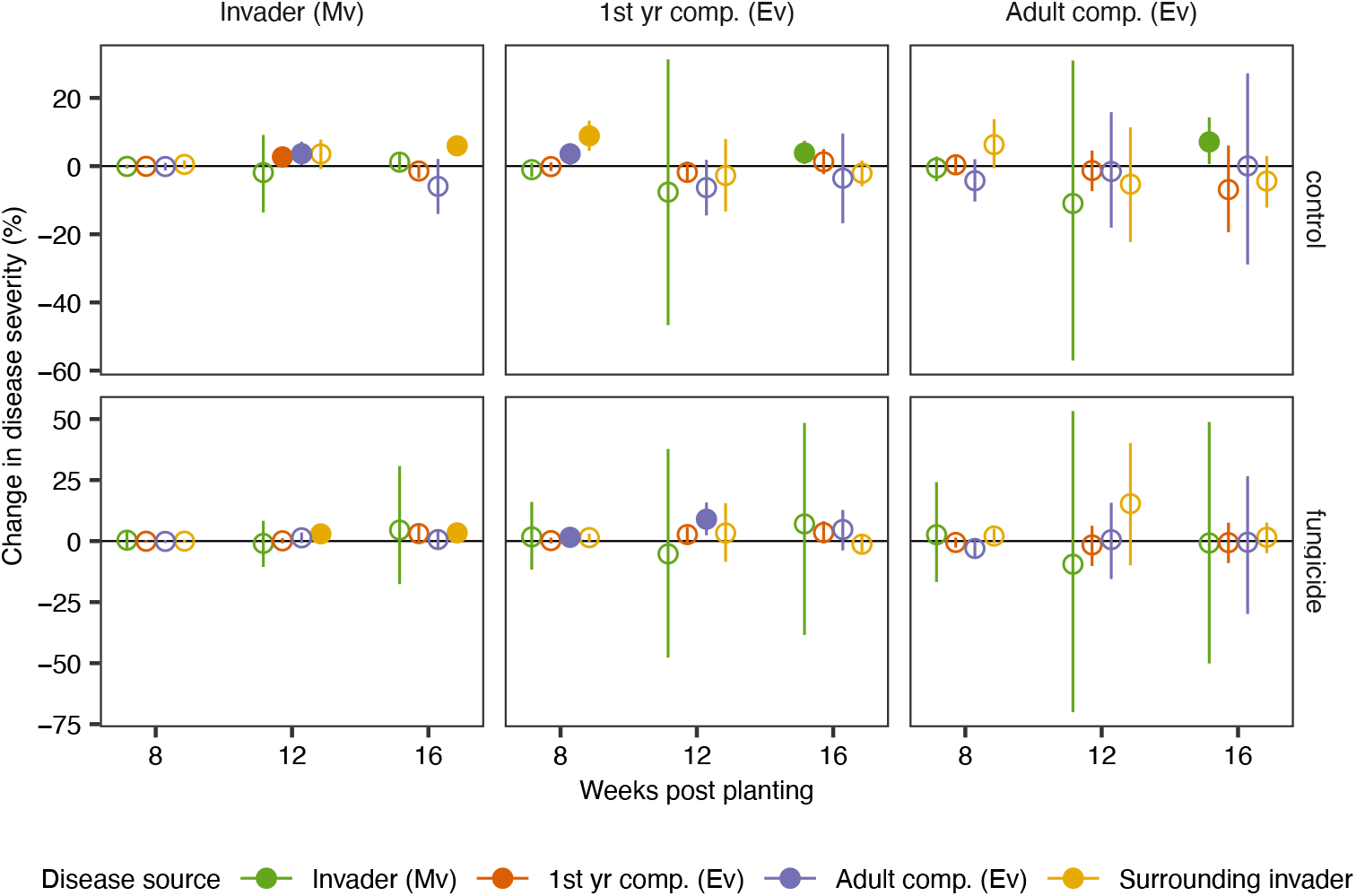
Disease transmission throughout the growing season, measured as change in disease severity (percentage of leaf area covered by lesions) on the invader, adult competitor, and first-year competitor (panel columns) with one unit increase in disease source four weeks prior. Disease sources included the density-scaled disease severity of invader, first-year competitor, and adult competitor plants, as well as disease severity of invader plants surrounding plots. One unit of density-scaled disease severity is one plant with 100% disease severity or 100 plants with 1% disease severity and one unit of surrounding invader (which occurred at high density) is an average disease severity of 1%. Model-estimated mean and 95% credible intervals are shown. Filled points have 95% credible intervals that omit zero.

Disease negatively affected key aspects of invader and competitor fitness. Relative to control plots, fungicide reduced disease severity on invaders by 41% [-60%, -18%], first-year competitors by 53% [-68%, -33%], and adult competitors by 21% [-55%, 23%] (Fig. 4A, Table S10). Reducing disease decreased competitor germination by 25% [-41%, -7%], but did not affect invader germination (Fig. 4B, Tables S15–S16). However, seed fungal infections, which were more common in control plots (Fig. S3A, Table S17), decreased invader germination by up to 44% [-80%, -11%] (Fig. S3B, Table S18). Reducing disease increased competitor establishment by 5% [0.4%, 12%] and did not change invader establishment, which was already 99% (Fig. 4C, Table S19). Unexpectedly, reducing disease decreased adult competitor biomass and seed production by 35% [-65%, -0.6%] and 26% [-47%, -2%] (Fig. 4D–E, Tables S4, S6), respectively, despite absence of a direct fungicide effect in the greenhouse (Methods S2, Tables S20–S22). When interactions were measured across biomass gradients (in contrast to density, as in Fig. 2A), reducing disease consistently increased adult competitor tolerance of competition (Fig. 4F– H, Fig. S4–S5), although the changes were not significantly different from zero (Table S23).

**Figure 4.**
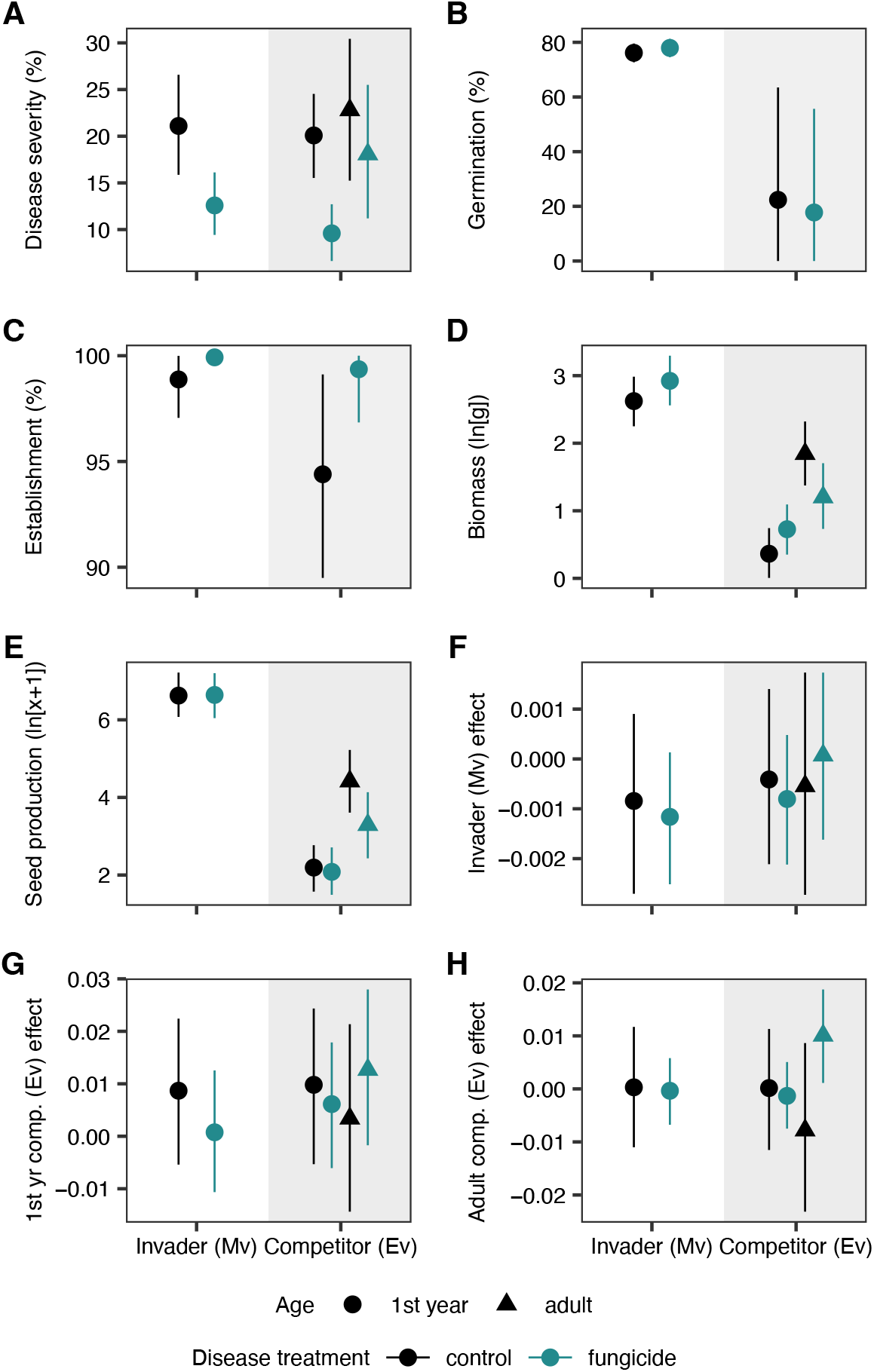
Effects of foliar fungal disease suppression on the invader (Mv) and competitor (Ev), measured as (A) disease severity, (B) germination (i.e., percentage of seeds that germinated in lab-based trials), (C) establishment (i.e., percentage of seedlings that established and survived through the growing season), (D) biomass (natural log-transformed), (E) seed production (natural log-transformed), and effects of (F) invaders, (G) first-year competitors, and (H) adult competitors (natural log-transformed change in focal plant biomass per unit background biomass). Model-estimated mean and 95% credible intervals are shown. Background shading delineates invader (white) and competitor (gray) values.

These processes, when put together in a model of long-term dynamics, predict that disease fails to mitigate invader impact, and instead enhances invader impact. Invasion reduced competitor biomass by 8% in the simulation (Fig. 5A), indicating limited competitive impacts of the invader in the absence of disease (Fig. S6). Disease emergence further decreased competitor biomass by 88%, and reduced invader biomass by 18%. Community-wide disease reached 50% (Fig. 5A). In contrast, the competitor did not maintain disease without the invader and its biomass remained unchanged (Fig. S7). When invader germination was reduced in the model due to seed infections (−44%), community-wide disease severity periodically fell below the threshold needed for disease effects (15%), leading to spikes in invader and competitor abundances. However, impacts on competitor biomass remained otherwise consistent (Fig. S8). Overall, the effects of disease on the invader measured in our experiment did not suppress it enough to allow the competitor to recover to its pre-invasion biomass.

**Figure 5.**
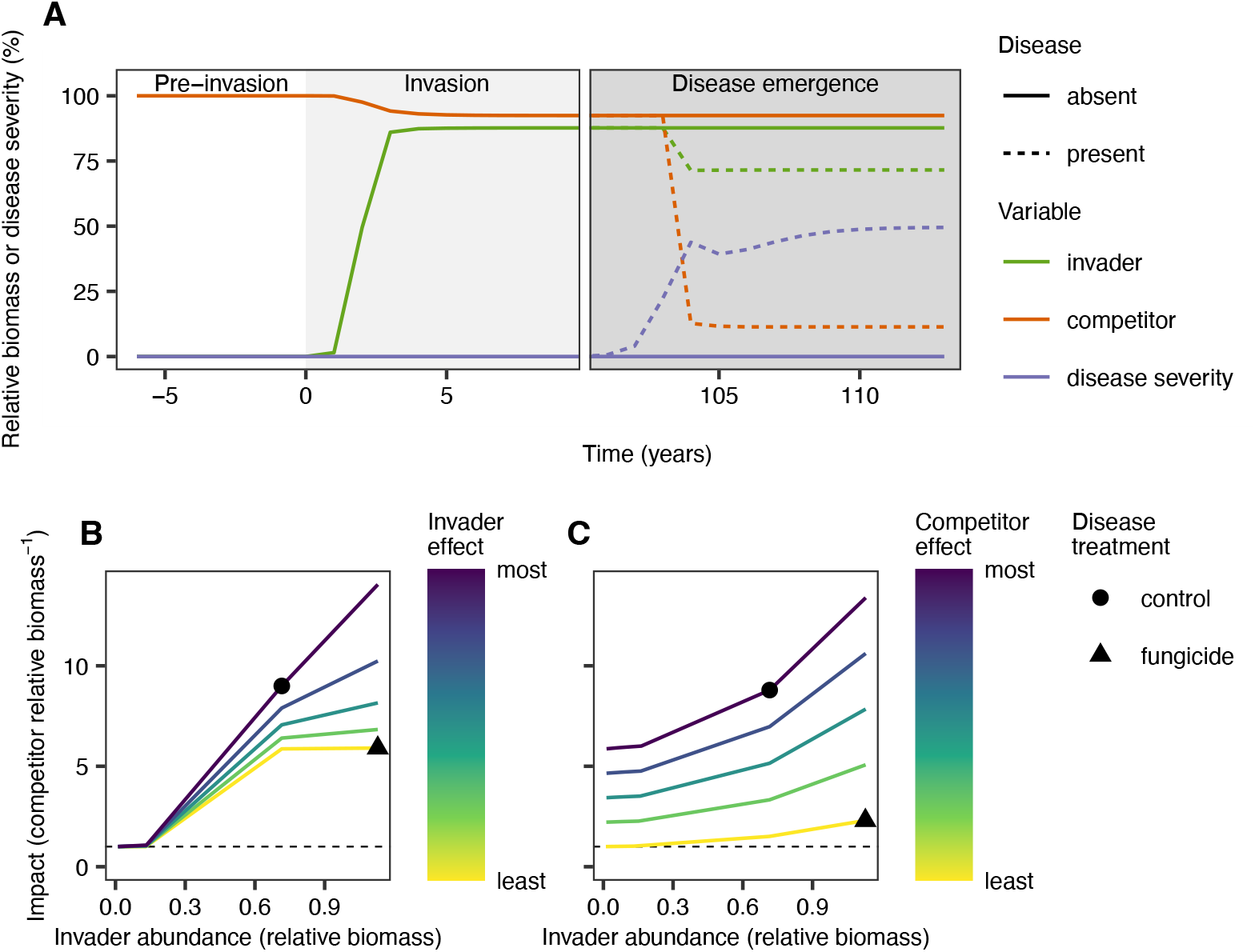
Simulations of population dynamics of the invader and competitor. (A) In a time series with parameter values based on field measurements, the invader reduced competitor abundance upon invasion and further reduced competitor abundance when disease emerged (years between 10 and 100 are omitted for visualization). Relative biomass is biomass divided by the biomass of the competitor pre-invasion. (B) Decreasing invader abundance and effects reduced the impact on the competitor, allowing it to recover to pre-invasion biomass at low invader abundance (dashed line). Parameter ranges: invader intraspecific effect = 0.001–0.09, invader interspecific effect = 0–5.4 × 10^−4^. (C) Reducing the competitor’s intraspecific effect led to competitor recovery. Parameter ranges: invader intraspecific effect = 0.001–0.09, adult competitor intraspecific effect = 0.001–0.008. In B and C, points are parameter values estimated from field measurements.

We evaluated the influence of invader abundance and per capita effects on impact (i.e., change in competitor biomass relative to its pre-invasion level) by varying competition coefficients. Because competition coefficients were based on biomass (Fig. 4F–H), with some modifications (see Methods), “per capita” refers to per g/m^2^. Disease decreased invader abundance, but increased invader effects, leading to greater impact (Fig. 5B). Similarly, disease increased the competitor’s per capita effect on itself, leading to greater impact (Fig. 5C). However, decreasing the competitor’s effect allowed the competitor to recover more than decreasing the invader’s effect (Fig. 5B–C). Other parameter settings that reduced invader abundance also allowed the competitor to recover (Fig. S9).

## Discussion

Enemy accumulation on invasive species is widespread (9, 10), but consequences for invader impacts are poorly understood (11). We demonstrated that recently emerged disease on the invasive annual grass *M. vimineum* suppressed both the invader and a perennial native competitor, *E. virginicus*, but the competitor was suppressed more. It is unclear whether the fungal pathogens (species in the genus *Bipolaris*) established in *M. vimineum*-invaded habitats before or after invasion. The first description of *Bipolaris* on *M. vimineum* in the U.S. was in 2001 (31), nearly a century after the first U.S. records of *M. vimineum* (32) and *Bipolaris gigantea* (33), suggesting that *Bipolaris* was maintained by resident species before aquiring the necessary tools to infect the invader. Invader amplification of *Bipolaris* is therefore likely a case of “spillback.” We found that enemy accumulation by an invader enhanced invader impacts through spillback and apparent competition.

Host-pathogen interactions can vary substantially over time and space. Plot-level environmental conditions did not explain variation in disease severity in our experiments, which may be due to limited environmental variation at this scale compared to larger spatial and temporal scales. For example, prior research on *M. vimineum* has found more intense impacts of disease in other locations (24, 25) and at the same location in previous years (29). Even when we assumed that disease had stronger effects on the invader in our model, the competitor did not recover to its pre-invasion biomass because disease increased its own intraspecific competition coefficient. This result depends on the pathogen’s ability to infect both species, whereas specialist pathogens accumulated by invaders may be expected to lessen invader impacts (12, 13). We estimated small impacts of the invader on the competitor in the absence of disease. Because plants were grown separately prior to planting in the field, we may have missed important competitive interactions among seedlings (34). However, the result that invader-amplified disease negatively impacted the competitor holds regardless of pre-disease competition.

Our results suggest that the invader is a more competent host than the competitor, sustaining disease transmission later in the growing season and throughout the simulation. However, the competitor suffered greater negative effects of disease. One possible explanation is asymmetric and inverse relationships between susceptibility (disease impact on biomass) and competency (inoculum production) of the hosts, as has been described for hosts of sudden oak death in California forest ecosystems (35). In addition, other native grasses outside the purview of this study may serve as reservoir hosts on which *Bipolaris* has historically perennated without the invader. Interestingly, fungicide decreased biomass, seed production, and germination of competitors without affecting its growth in the greenhouse, suggesting a potential negative effect of fungicide on beneficial fungi in the field.

The effects of enemy accumulation on invader impacts have been evaluated in few systems, with a range of outcomes. For example, the invasive grass *Ammophila arenaria* increased seed predation of a native legume by a native rodent, which reduced the predicted growth rate of the legume (36). In contrast, a vascular wilt disease acquired by *Ailanthus altissima* in its invaded range seemed to have limited effects on most co-occurring species (37, 38). In a greenhouse study, we found that *Bipolaris* inoculation decreased biomass of *E. virginicus*, but not two other native species (34). A general rule for how enemy accumulation by invaders affects competitors is therefore difficult. Instead, host screening, as in biological control development (38), paired with field studies and population modeling (39), can increase understanding of how enemy accumulation will affect native competitors and inform management strategies.

Mitigating invader impacts is important for maintaining and restoring ecosystems. Increasing documentation of new enemies associated with invaders (9, 10), and lessons from research on biocontrol (12, 13), the enemy release hypothesis (6), and biotic resistance (40) have led to the hypothesis that enemies may mitigate invader impacts over time (8). This hypothesis is relevant to a broad range of invasive taxa that share enemies with co-occurring species (20, 41, 42). By pairing field experiments manipulating plant density and disease with a dynamical model, we demonstrated that although accumulated enemies may cause invader declines, they can also increase invader impacts because of apparent competition. We expect this result to apply broadly because most enemies accumulated by invaders are generalists (to be present pre-invasion, enemies must be sustained by alternative hosts, and so are unlikely to be specialists) and invaders often reach high abundances (4, 5), which can amplify enemies. Continued research can illuminate whether the effects of spillback persist over evolutionary timescales as enemies adapt to invaders.

## Materials and Methods

### Density experiment

To evaluate the effects of disease on the invader abundance–impact relationship, we carried out a two-year field experiment at Big Oaks National Wildlife Refuge (BONWR) in Madison, IN, USA from April 2018 to October 2019. Invader (*M. vimineum*) populations are established in approximately 810 ha of BONWR’s 20,647 ha property, and some are infected with *Bipolaris* fungi (25, 29, 30). The experiment was a full factorial combination of two disease treatments (fungicide, control) and ten density treatments, replicated four times each—once at each of four sites with the invader and *Bipolaris* present (Fig. S10).

In spring 2018, we planted focal plants— three invader seedlings, three first-year competitors, and one adult competitor—in the center 1-m^2^ of each 2 m × 2 m plot (Fig. S11). We also planted density treatments in the center of plots, including: no additional plants or low, medium, or high densities of the invader (4, 16, or 64 plants), first-year competitors (4, 8, or 16 plants), or adult competitors (2, 4, or 8 plants). Invaders, first-year competitors, and dead adult competitors were replaced in spring 2019. Every four weeks during the growing season, we sprayed fungicide-treated plots with 1.5% fungicide (Ipro 2SE, 23.8% iprodione) and control plots with tap water, which did not promote disease relative to ambient levels (Table S24).

We monitored focal plants for survival and foliar fungal infection throughout the growing season and measured growth and seed production at the end of the growing season. We measured seed germination and, for invaders, seed fungal infection, in the lab. We also measured environmental variables, including soil moisture, canopy cover, temperature, and relative humidity in each plot. Due to changes in data collection methods between years, results are presented from the second year unless otherwise noted. Additional details are described in Methods S1 and Table S25.

### Litter experiments

We conducted two field experiments to evaluate effects of disease and invader litter on invader and competitor emergence. In spring 2018, we established seven treatments, replicated once at each of four sites at BONWR (Fig. S10, Methods S3). The invader was established at these sites and showed no evidence of foliar fungal infections. We collected invader litter with evidence of *Bipolaris* infections from elsewhere at BONWR. We autoclaved half of this litter for 60 minutes and brought all back to the field. We cleared 2 m × 2 m plots and sowed 200 invader seeds and 50 competitor seeds in the center 1-m^2^. Plots received either no litter, autoclaved litter, or control litter. The autoclaved and control litter were each applied at low (50 g/m^2^), medium (100 g/m^2^), and high (200 g/m^2^) levels. We measured seedling emergence in July. We were unable, however, to identify competitor seedlings amongst other species in the seedbank.

Because we could not estimate competitor emergence from the first litter experiment, and because there was no significant effect of autoclaving on invader emergence (Fig. 1C), we performed a second litter experiment to estimate competitor germination with non-autoclaved litter. In April 2019, we established three litter treatments—removal, addition, and unmanipulated—replicated four times (two replicates each near sites D1 and D3; Fig. S10). We raked and weighed litter from each 2 m × 2 m plot, adding litter from removal plots to addition plots. Prior to returning litter to plots, we planted 50 competitor seeds in removal plots and 26 competitor seeds in addition and control plots. Seeds were purchased from Prairie Moon Nursery and glued to toothpicks to aid seedling identification. We measured tiller emergence in June.

### Statistical analyses

We fit Bayesian linear regressions using the *brms* package (43) in R (version 4.0.2) (44). Tables S1–S24 describe model formulae, prior and posterior distributions, and model estimates. We used the concept of constant final yield (i.e., that plant biomass saturates with increasing density) to estimate relationships between invader abundance and density (Fig. 2A) and a Ricker model to estimate competition coefficients (Fig. 2B, 4F–H). Models were run with three chains of 6000 iterations each with a discarded warm-up period of 1000 iterations. We evaluated convergence and fit by plotting simulated data from the posterior predictive distributions against observed data, ensuring the three chains mixed well, and checking that r-hat values equaled one. We used the hypothesis (43) and emtrends (for effects of continuous variables on disease severity) (45) functions to extract estimates and 95% credible intervals of values of interest.

### Dynamical model

We simulated the long-term population dynamics of the invader and competitor using a continuous-time growing season model linked to a discrete-time yearly model (Methods S7). In the continuous-time model, we tracked the dynamics of plant biomass, which changes as plants grow, compete, and transmit disease during the growing season. The discrete-time model projects the dynamics of adult competitors and seeds of both species at the beginning of the growing season each year, which are determined at the end of the continuous-time model by survival and seed production, respectively. Seed number is determined by seed production the prior year and carry-over of seeds in the seed bank. Seed germination and establishment occur quickly at the start of the growing season, and so are included as part of the discrete-time yearly model. The density of newly established plants then initiate biomass at the beginning of the continuous-time growing season model. Parameters derived from the density experiment were set to values estimated from control plots when community-wide disease severity exceeded 15% and fungicide-treated plots otherwise.

We performed simulations in R using the *deSolve* package (46). We began simulations with ten competitor seeds. After 100 simulated years, we introduced ten invader seeds. After 100 more years, we introduced fungal inoculum. We assumed the growing season lasted 160 days. Competition coefficients from Fig. 4F–H were used, except that the effect of fungicide on the invader’s intraspecific coefficient was estimated from plot-scale biomass (Fig. 2A) and positive coefficients (i.e., facilitation) were set to zero. However, because adult competitors then exerted no intraspecific competition on adults in fungicide plots and first-year competitors in control plots, we replaced those estimates with estimates from adult competitor effects on first-year competitors in fungicide plots and adults in control plots, respectively.

## Acknowledgments

We thank Kristin Baecher, Liliana Benitez, Zobia Chanda, Taylor Clark, Laney Davidson, Katelyn Doyle, Riley Dunlop, Trevor Green, Mariam Higginbotham, Zadok Jollie, Evan Lacey, Daniela Menendez, David Notman, Teresa Orosa, Shannon Regan, Penny Reif, Callie San Antonio, Vida Svahnström, Thomas Thrasher, Ryan Truesdell, and Max Zaret for assistance with the study. We thank Joe Robb and the staff of Big Oaks National Wildlife Refuge for site access and assistance. We received valuable feedback on the manuscript from members of the Flory and Holt labs. Fig. 1 created with BioRender.com. This work was funded by USDA award number 2017-67013-26870 as part of the joint USDA-NSF-NIH Ecology and Evolution of Infectious Diseases program.

