## Supplementary Information for "Effects of disease emergence on invasive grass impacts"

### Supplementary Information for
#### Effects of disease emergence on invasive grass impacts

Amy E. Kendig, Ashish Adhikari, Brett R. Lane, Christopher M. Wojan, Nicholas Kortessis, Margaret Simon, Michael Barfield, Philip F. Harmon, Robert D. Holt, Keith Clay, Erica M. Goss, and S. Luke Flory

**Methods S1.** Additional methods for density experiment

*Experimental set-up*

We selected four sites at BONWR that contained *M. vimineum* populations with evidence of previous *Bipolaris* infection (i.e., we observed characteristic lesions on litter in April 2018). We established 20 2 m × 2 m plots at each site, avoiding mature tree and shrub stems, and haphazardly assigned an experimental treatment to each one. At each site, plots occupied approximately 630–1350 $m^2$ and plots centers were at least 2.8 m from one another. Each April, before invader emergence, we removed plant material from plots by raking leaf litter and spraying live plants with 4% glyphosate herbicide (GlyStar Plus, 41% glyphosate).

We grew invader and competitor (*E. virginicus*) seedlings in a greenhouse at Indiana University (Bloomington, IN, USA). These included "focal plants" from which we measured responses to the treatments and "background plants" that form the density treatments. Seeds were collected from BONWR in 2017 and, for invader only, 2018. Seedlings grew for five weeks (except competitors in 2018, which grew nine weeks) in potting soil (peat/Sungro Metro-Mix, vermiculite, perlite, and osmocote in a ratio of 2:1:1:1) with ambient conditions. Seedlings were placed outside of the greenhouse for acclimation approximately one week prior to planting in the field (Table S25).

Seedlings were planted in early May, and in 2018, we also transplanted adult competitor plants from a nearby site at BONWR into plots. In spring 2019, surviving adult competitors were left in the plots while all other plants were replaced.  The outer 0.5 m perimeter of the 2 m × 2 m plot was left without plants to buffer against external plant and litter encroachment. The competitor density gradients spanned natural *E. virginicus* densities at our site, which reached approximately 2.3 $m^{-2}$ in the presence of the invader (Methods S4). The invader density gradient was within the range of natural densities at our site, which can reach over 560 $m^{-2}$ (1).

Plots were sprayed with 1.5% fungicide solution or tap water using backpack sprayers until the liquid dripped off the leaves once every four weeks from planting in May to late September (Table S25). The fungicide Iprodione was developed to kill some foliar fungi associated with turfgrass and ornamental plants and is effective against *Bipolaris* infection (2). Based on a greenhouse experiment, there was no significant direct effect of iprodione on invader biomass, seed heads, or competitor biomass (Tables S20–S22, Methods S2).

In 2019, we adhered to a strict weeding schedule to maintain the planted densities (Table S25), but this was not the case in 2018. Invaders from the seedbank and outside of the plots grew into plots in 2018. Starting in July 2018, we weeded encroaching invader plants from the plot edges, but did not remove them from inside of the plot to avoid accidentally removing experimental plants.

Plants that died were replaced in May in 2018 and throughout the experiment in 2019. In 2018, a tree fell on two plots in site D4 (fungicide-treated plots planted with medium and high densities of adult competitors), so they were removed from analyses for that year. The plots were added again in spring 2019 in new, nearby locations.

*Data collection*

<u>Survival</u>— We recorded focal plant mortality every four weeks throughout the growing season. Plants with no green leaves and no evidence of seed production were recorded as dead. To

maintain density treatments, dead plants were immediately replaced in 2019; however, we did not replace them in 2018. We estimated winter survival of competitor plants in April 2019 by recording mortality of plants that were alive in September 2018.

Growth—We counted the number of tillers for all focal plants in June and July 2018. Invader tiller numbers increase over the growing season in proportion to biomass (3). During October 2018, we clipped aboveground invader biomass within a 0.25 × 0.5 m quadrat placed over the center focal invader in each plot. During October 2019, we harvested all aboveground invader and competitor biomass by cutting plants at the soil surface. The invader has a mixed mating system, including obligately-selfed cleistogamous seeds located within leaf sheaths and potentially outcrossed and exposed chasmogamous seeds (4). In both years, we separated cleistogamous seeds from invader biomass and oven-dried all biomass at 60°C until constant weight (48–72 hr).

Seeds—In September of both years, we placed 9.5 cm × 12.7 cm white organza drawstring bags on invader chasmogamous seed heads (ten seed heads per plot in 2018 and five seed heads per focal plant in 2019). We collected the bags four weeks later and counted the number of seeds per bag. In 2018, we estimated seeds per plant by multiplying the July tiller counts by the average number of seeds per seed head for the same plot. In 2019, we counted the number of chasmogamous seed stems— terminal stems without leaf sheaths—per focal plant and multiplied this number by the average number of seeds per seed head for the same plant. To collect cleistogamous seeds in 2019, we crumpled invader biomass over a series of sieves, removed non-seeds, and weighed the seeds. We fit a linear regression to cleistogamous seeds per plant for 15 plants from a range of sites and treatments (seeds = 68.22 + 1079.43 × seed weight in grams, adjusted $R^2 = 0.99$, $p < 0.001$). We also measured cleistogamous seeds in 2018 but omitted these from analyses as they could not be converted to an amount per plant.

We collected spikelets as they ripened from focal competitor plants throughout the growing seasons of both years. We dissected spikelets from 36 plants from a range of sites, treatments, and ages (i.e., adult and first-year) across both years and fit a linear regression to convert spikelet weights from all plants to seed numbers (seeds = 72.798 × spikelet weight in grams, adjusted $R^2 = 0.96$, $p < 0.001$).

Germination and seed infection—We surface-sterilized chasmogamous invader seeds collected in 2018 with a one-minute wash of 20% household bleach. We rinsed seeds four times with sterile deionized water and plated them on sterile antibacterial agar plates. All supplies were sterilized by autoclave or 20% household bleach. We sealed the plates and stored them in a growth chamber with a 12:12 h light:dark cycle at 23.8ºC:15.5ºC for ten days. We counted the number of seeds with fungi growing from them, dividing counts by fungi with light hyphae and fungi with dark hyphae. Morphological characterization of a subset of fungi growing from invader seeds suggested that dark-hyphae fungal infections include species in the genera *Bipolaris*, *Pyrenophora*, and *Dreschlera*, and light-hyphae fungal infections also include *Bipolaris* species. We also counted the number of seeds that germinated. Three replicate petri dishes were prepared for each plot.

To estimate competitor seed germination, we planted up to thirty seeds per age–plot combination from plots with no background plants, high invader density, high first-year competitor density, or high adult competitor density in both years. Seeds were planted in growing medium and trays in a greenhouse at Indiana University (as described in *Experiment set-up*) and watered twice daily. Seedlings were counted after two, three, and four weeks and seeds were sieved and examined after four weeks. We combined the data from both years when evaluating the effect of fungicide on germination (Table S16). We did not evaluate competitor seeds for fungal infection.

Foliar fungal infection—In July, late August, and September of 2018 and every four weeks between June and late August in 2019, we counted the total number of leaves and the number of leaves with at least two foliar lesions on a haphazardly selected tiller of each focal plant and we collected a leaf with a moderate amount of foliar lesions for the plant. We scanned the collected leaves and used ImageJ (version 2.0.0) (5) to calculate the leaf area and proportion of leaf area covered by brown lesions. The area classified as lesions contained some senescing plant tissue that may or may not have been the result of fungal pathogen infection. We reviewed each output image and edited scans to correct for poor estimation of lesion area.

To estimate disease severity (percentage of tissue with lesions), we divided the leaf area with lesions per tiller by the total leaf area per tiller. To estimate sources of disease, we divided the average leaf area with lesions per tiller by the average total leaf area per tiller for a given plant group (invaders, first-year competitors, adult competitors) in each plot, multiplied the proportion by the planted density of the plant group in the plot. When evaluating the effect of fungicide on overall disease severity (Fig. 4A), we used data from late August for the invader and early August for the competitor—when disease severity was high and the fungicide effect was most obvious. In July, August, and September of 2018 and July and August of 2019, we collected leaves from each species in each plot and examined them for conidiophores characteristic of *Bipolaris* fungi using microscopy (6).

To characterize disease on invader plants surrounding experimental plots, we collected invader biomass outside each plot with a circular subplot in July 2018. We separated leaves and stems into healthy or infected, oven-dried them, and weighed them. We used the proportion of biomass that was infected as a proxy for surrounding disease severity. In 2019, we collected 20 haphazardly selected invader leaves from around the outside of each plot each month. We scanned the leaves, quantified leaf and lesion areas, and calculated disease severity as the percentage of leaf area with lesions across all 20 leaves.

<u>Environmental variables</u>—We measured soil moisture with a Hydrosense II (Campbell Scientific, Inc.) in each plot in June 2018. In July 2018, we measured canopy cover above each plot with a densiometer. From July to October 2019, we measured temperature and humidity in the center of each control plot with HOBO U23 Pro v2 data loggers (Onset Computer Corporation). We estimated average monthly dew intensity by multiplying the daily difference in dewpoints when the air was saturated (relative humidity = 100%) by the number of hours the air was saturated and averaging the product across days in a month. We used dew intensity as a proxy for leaf wetness duration, which promotes disease severity caused by *Bipolaris gigantea* (7).

**Methods S2.** Direct effects of iprodione fungicide

To evaluate the effects of iprodione fungicide on plant growth in the absence of *Bipolaris* foliar fungal infection, we grew the invader (*Microstegium vimineum*) and competitor (*Elymus virginicus*) in 20 pots each in a greenhouse at the University of Florida (Gainesville, FL, USA). We collected invader seeds from BONWR in fall 2015 and purchased competitor seeds from Prairie Moon Nursery (Winona, MN, USA) in spring 2018 and stored them at 4℃ prior to the experiment. On February 28, 2019, we planted seeds in Metromix 930 growing medium (Sungro Horticulture) saturated with tap water.

Seedlings grew for eight weeks, during which time we transplanted them from seed starting trays to 1 L pots and, to simulate methods of the field experiment (Methods S1), placed them outside in a shaded area for three days before returning them to the greenhouse. Then we applied a fungicide treatment consisting of 1.5% Ipro 2SE (23.8% iprodione, ADAMA) dissolved in deionized water to half of the pots and deionized water as a control to the other half. We sprayed plants until liquid dripped off the leaves (approximately 137 ml per plant). We re-applied the treatment once for the invader, which was harvested six weeks after treatments began because it began producing seeds. We re-applied the treatments to the competitor every four weeks for 30 weeks. Plants were watered daily with a drip irrigation system and haphazardly rearranged on greenhouse benches twice per week. Garden Safe insecticidal soap was occasionally applied to all plants to help control aphids and thrips.

We harvested the competitor 34 weeks after treatments began. We counted the number of seed heads per plant for the invader (only one competitor plant produced seeds) and weighed oven-dried biomass (dried to constant mass) for both species. Final sample sizes were 9 plants per species for the control treatment and 10 plants per species for the fungicide treatment.

**Methods S3.** Additional methods for first litter experiment

*Experimental set-up*

In April 2018, we removed the plant material from 13 2 m $\times$ 2 m plots at each of four sites at BONWR (Fig. S10), by raking away leaf litter and spraying live plants with 4% glyphosate herbicide (GlyStar Plus, 41% glyphosate). On May 9, we collected invader litter with visible signs of foliar *Bipolaris* infection from BONWR. We brought the litter to Indiana University in Bloomington, IN, and prepared 112 50-g aliquots for low (50 g/m$^2$), medium (100 g/m$^2$), and high (200 g/m$^2$) litter levels. The high weight is representative of litter amounts found in a heavily invaded area of BONWR (Methods S5). Half of the aliquots were autoclaved in loosely closed autoclave bags for 60 minutes and then transferred to paper bags. A pilot experiment demonstrated negligible effects of autoclaving on litter weight (Methods S6).

The full experiment included 13 treatments and only the inner 1-m$^2$ of plots received treatments. The outer 0.5 m perimeter buffered against external plant and litter encroachment. Plots received either no litter, autoclaved litter, or control litter. On May 11, the autoclaved and control litter were each applied at low, medium, and high levels. The litter treatments were repeated twice: once with plants and once without plants. For plots with plants, we added 200 invader seeds in 1/3, 50 competitor seeds in 1/3, and a single adult competitor plant in 1/3 (all prior to adding litter). Seeds were collected from BONWR in the fall of 2017 and adult competitor plants were transplanted from a nearby BONWR site with no evidence of foliar fungal infection on the invader. The treatments without plants were controls designed to quantify the contribution of invader seeds from the litter.

*Data collection*

On July 6, we counted the number of invader seedlings that had emerged in the third of each plot where they were sowed. Counts of competitor seedlings were not reliable due to the difficulty in identifying single-blade grasses down to species. We used the third of the plots in which competitor seeds were sowed to estimate the number of invader seedlings coming from the seedbank or litter. While the plants without litter were meant to serve as these controls, low emergence of competitor seeds meant that the 1/3 of the plot where they were planted was relatively free of plants other than the invader. In addition, we assumed the sub-plot next to the invader's subplot was a better estimate of the seedbank than a plot further away.

**Methods S4.** Measuring natural densities of competitor (*E. virginicus*)

On June 7, 2018, we placed a 1 m × 1 m quadrat in three locations near the experimental sites at BONWR (Fig. S10) with both competitor and invader plants present, aiming to capture maximum densities of the competitor (invader densities were relatively high). Competitors were at least one year old and had produced seeds and invaders were seedlings. We counted the number of competitor plants and found 1, 3, and 3 plants in the three locations (mean = 2.3 $m^{-2}$).

**Methods S5.** Determining litter quantities for first litter experiment

On April 4, 2018, we collected litter from six haphazardly selected 0.5 m $\times$ 0.5 m plots at the BONWR site described in the main text. We cut the litter around the inside edges of the PVC sampling frame used to delineate the plots and put it in plastic bags. The litter samples were then weighed at the University of Indiana (Bloomington, IN, USA). We did not separate invader litter from other types of litter, but invader appeared to be the dominant species. The litter weights were as follows:

| Plot replicate | Litter weight (g/m$^2$) |
|---|---|
| A | 197.36 |
| B | 291.00 |
| C | 369.76 |
| D | 364.76 |
| E | 458.88 |
| F | 204.72 |
| Average | 314.41 |
| Standard error | 41.93 |

**Methods S6.** Evaluating the effect of autoclaving on litter weight

In April 2018, we prepared six aliquots of litter using the material described in Methods S3. These aliquots were individually weighed and placed in labelled autoclave bags, which were loosely rolled and taped. All bags were autoclaved for 60 minutes. Litter was then removed from the bags and weighed again. Based on these data, we determined that autoclaving has negligible effects on the weight of litter.

| Replicate | Pre-autoclave weight (g) | Post-autoclave weight (g) | Percent change |
|---|---|---|---|
| 1 | 100.75 | 98.33 | -2.40 |
| 2 | 106.36 | 101.711 | -4.37 |
| 3 | 49.82 | 47.31 | -5.04 |
| 4 | 53.21 | 50.56 | -4.98 |
| 5 | 157.78 | 151.54 | -3.95 |
| 6 | 154.19 | 146.82 | -4.78 |
| Average | | | -4.25 |

**Methods S7.** Additional methods for the dynamical model

*Growing season dynamics*

For the continuous-time growing season model, time is measured in days ($\tau$). We track both the total aboveground biomass density ($B_i$) and infected biomass density ($I_i$) of the invader (annual plant, $i = A$), adult competitors (perennial plant, $i = P$), and first-year competitors ($i = F$). Because the invader produces large amounts of litter that can harbor fungal pathogen spores (2, 8), we also model biomass density of newly produced litter ($D$) and biomass density of litter-derived fungal conidia ($C$). Each component changes during the growing season according to

$$\frac{dB_i}{d\tau} = r_i B_i(1 - \alpha_{iA}B_A - \alpha_{iF}B_F - \alpha_{iP}B_P) - m_i B_i - v_i I_i \tag{3}$$

$$\frac{dI_i}{d\tau} = S_i(\beta_{iC}C + \beta_{iA}I_A + \beta_{iF}I_F + \beta_{iP}I_P) - (m_i + v_i)I_i$$

$$\frac{dD}{d\tau} = m_A B_A + v_A I_A + m_F B_F + v_F I_F + m_P B_P + v_P I_P$$

$$\frac{dC}{d\tau} = h\big((m_A + v_A)I_A + (m_F + v_F)I_F + (m_P + v_P)I_P\big) - aC$$

$$S_i = B_i - I_i$$

where $S_i$ is the susceptible biomass of plant group $i$ ($i = A$, $F$, or $P$). Total biomass grows with intrinsic growth rate $r_i$, is inhibited by competition from each plant group, and is lost through senescence or shedding at rate $m_i$. We assume susceptible and infected tissue both contribute to biomass growth. Infected biomass increases with density-dependent transmission and decreases through $m_i$ and an additional loss term for virulence, $v_i$. All dead biomass becomes litter. Conidia in the litter come from dead infected biomass at rate $h$ and decline at rate $a$. Newly produced litter does not affect dynamics during the growing season (except through production of $C$), but reduces seedling establishment at the start of the following growing season (see Eqs. 4–5). Initial values at the start of the growing season are determined by the discrete-time model of yearly dynamics (described below).

*Yearly dynamics*

For the discrete-time yearly model, time is measured in years ($t$). Building upon previous models of annual and perennial plant dynamics (9, 10), we assume yearly population dynamics change according to

$$A[t+1] = s_A(1 - g_A)A[t] + c_A B_{A,T}[t] \tag{4}$$

$$F[t+1] = s_P(1 - g_P)F[t] + c_F B_{F,T}[t] + c_P B_{P,T}[t]$$

$$P[t+1] = l_P P[t] + g_P E_P[t] l_P F[t]$$

$$L[t+1] = B_{A,T}[t] + B_{F,T}[t] + B_{P,T}[t] + D_T[t] + (1 - d)L[t]$$

where $A$ is the density of invader seeds, $F$ is the density of competitor seeds, and $P$ is adult competitor density present at the beginning of the growing season. Seeds at this time come from two sources: seed production the prior year and between-year survival in the seed bank, which is represented by no germination (1-$g_i$) and seed survival ($s_i$). Each plant group $i$ produces $c_i$ seeds per unit end-season biomass, such that newly produced seed is $c_i B_{i,T}[t]$, where the subscript $T$ indicates $\tau$ is equal to the final day of the growing season. Adult competitors survive through the year at rate $l_P$ or are produced when seeds germinate ($g_P$), establish ($E_P$), and survive to the next growing season ($l_P$). Finally, litter ($L$) is produced by live biomass at the end of the growing season and dead biomass generated during the growing season. Litter is lost between years with decomposition rate *d*. We assume litter decomposition begins the first summer following litter production. Litter can reduce the establishment of invader and competitors seeds, which is represented by

$$E_i[t] = \frac{e_i}{1+\gamma_i L[t]} \quad (5)$$

where $e_i$ is the maximum establishment fraction of plant group *i* (*i* = *A* or *P*) in the absence of litter and $\gamma_i$ is the sensitivity of establishment to litter.

All plant biomasses at the beginning of the growing season are susceptible ($B_{i,0} = S_{i,0}$, where the subscript 0 indicates $\tau$ is equal to the first day of the growing season). The initial biomasses of each plant group at the start of the growing season in year *t* are

$$B_{A,0}[t] = S_{A,0}[t] = g_A E_A[t] b_A A[t] \quad (6)$$

$$B_{F,0}[t] = S_{F,0}[t] = g_P E_P[t] b_F F[t]$$

$$B_{P,0}[t] = S_{P,0}[t] = N_P[t]\big(B_{F,T}[t-1] + B_{P,T}[t-1]\big)$$

where $b_i$ is the initial biomass of a first-year individual of plant group *i* that germinates ($g_i$) and establishes ($E_i$). Adult competitors begin with new biomass that is proportional (through $N_P$) to surviving biomass at the end of the previous growing season. Here, we are assuming that larger plants produce larger belowground perennating organs, which promote growth in the following year (11). We also assume that litter limits the initial growth of perennials, and represent the proportional contribution of biomass from the previous year to new growth as

$$N_P[t] = \frac{n_P}{1+\gamma_P L[t]} \quad (7)$$

where $n_P$ is the maximum contribution of biomass from the previous year to new growth in the absence of litter and $\gamma_P$ is the sensitivity of biomass growth to litter. The initial conidia in the litter is the sum of conidia remaining from the previous growing season and new conidia produced from infected biomass at the end of the previous growing season

$$C_0[t] = C_T[t-1] + h\big(I_{A,T}[t-1] + I_{F,T}[t-1] + I_{P,T}[t-1]\big) \quad (8).$$

To include the effects of disease measured in the field experiment, we used parameter values from control plots (Table S26) when community-wide disease severity of live biomass at the end of the previous growing season exceeded 15% and parameters values from fungicide plots otherwise (Table S26). We calculate community-wide disease severity as

$$\big(I_{A,T} + I_{F,T} + I_{P,T}\big)/\big(B_{A,T} + B_{F,T} + B_{P,T}\big) \quad (9).$$

*Parameter estimation*

We estimated parameter values for the dynamical model using information from our experiment and prior studies (Table S26). We used estimates based on statistical models fit to data from our experiment or data from the experiment by Benitez et al. (12), as well as values from the literature.

To estimate the growth rate of plant group *i*, $r_i$, we used the intercepts and plant group coefficients from the statistical model fit to log-transformed biomass (Table S23, Fig. 4F–H). The intercepts represent the growth of each plant group in the absence of competition. Because we did not measure the initial biomass of plants in the field experiment, we assumed that plants began at 10% the minimum measured plant biomass (also the values assigned to $b_i$ for each plant group *i*).

We divided the change in biomass by 160 days as this was the approximate length of time plants were in the field. We also used this model to estimate the competitive effect of plant group *j* on group *i*, $\alpha_{ij}$.

To estimate transmission between plant groups *i* and *j*, $\beta_{ij}$, we used model-estimated disease severity changes (Table S10, Fig. 3). To estimate transmission from the invader, we chose the larger estimate between invader plants within plots and surrounding plots. For all transmission coefficients, we selected the largest positive estimate among all time points and set the value to zero if none existed. We multiplied estimates by $5 \times 10^{-4}$ to imitate observed disease severity values (as in Fig. 4A).

We estimated the conversion coefficients for biomass to seeds for each plant group *i*, $c_i$, using normal linear mixed-effects models fit to seeds per plant for each plant group. The regressions were fit with forced zero intercepts and the fixed effects were plant biomass and an interaction between fungicide and plant biomass. Plot was the random intercept for the invader and first-year competitor and site was the random intercept for adult competitors. We used normal distributions (location = 0, scale = 100) for the coefficient priors and half Student's *t*-distributions for the standard deviation priors (location = 0, df = 3, $\text{scale}_{\text{invader}}$ = 589.8, $\text{scale}_{\text{first-year competitor}}$ = 10.4, $\text{scale}_{\text{adult competitor}}$ = 70.3). The fungicide interaction was not significantly different from zero for any of the plant groups.

We estimated the maximum fraction of annual and perennial seedlings that establish in the absence of litter ($e_A$, $e_P$) with the statistical model fit to growing season survival data (Table S19, Fig. 4C). Survival of perennial adults over a year ($l_P$) was estimated with survival data recorded in April 2019 (plants were transplanted in May 2018). We used a Bernoulli (logit-link) generalized linear mixed-effects model with a random intercept for site. We used a normal distribution (location = 0, scale = 1) for the intercept prior and half Student's *t*-distributions (location = 0, scale = 2.5, df = 3) for the standard deviation priors. The fixed effect was fungicide treatment.

To estimate maximum contribution of the previous year's perennial biomass to new biomass, $n_P$, we calculated the proportion of competitor biomass that was alive at the end of the greenhouse experiment (Methods S2) and multiplied this by the fraction of adult perennials that survived over the year (described above). We also used data from the fungicide experiment to estimate biomass loss of perennials over the growing season, $m_F$, which we calculated as the proportion of biomass that senesced divided by 292 days (the length of the greenhouse experiment). Because we did not have comparable data for $m_P$ and $m_A$, we assumed they were equal to $m_P$.

To estimate litter sensitivity, $\gamma_A$ and $\gamma_P$, we used data from Benitez et al. (12). The design of this experiment was similar to our first litter experiment described in Methods S3, except we are able to get estimates for both the competitor and the invader. We fit Eq. 5 to the proportion of invader and competitor seeds that established over four litter amounts (0, 50, 100, and 200 g/m$^2$) for each species. The experiment contained 24 pots of 50 seeds each for each species. Plants grew in a greenhouse in Gainesville, FL over ten weeks. We used a non-linear function with a normal response distribution. We used a normal distribution (location = 0.85, scale = 1) for the $e_i$ prior, an exponential distribution (rate = 1) for the $\gamma_i$ prior, and a Cauchy distribution (location = 0, scale = 1) for the standard deviation prior. Benitez et al. (12) reported that 5500 fungal conidia were recovered per gram of invader litter, which we used as our estimate for *h*.

To estimate the litter decomposition fraction, we used data from the study by DeMeester and Richter (13). Briefly, three grams of standing *M. vimineum* litter were put in twelve 0.001 m$^2$ mesh bags (1 mm diameter openings) and placed in a forested riparian wetland in North Carolina, USA. The bags were placed in six plots dominated by *M. vimineum* and six plots where *M. vimineum* was periodically removed with hand-weeding. We extracted data collected after 360 days in Fig. 3 using the R packages "jpeg" and "zoo" (14–16). The average percentage of mass remaining in the two plot types was 41%, leading to an estimate of 0.59 for *d*. We applied this value to *E. virginicus* litter as well.

To estimate survival of perennial seeds that remain in the seed bank, $s_P$, we used data from the study by Garrison and Stier (17). Briefly, 100 seeds of *E. virginicus* were added to each of 30 polyester bags filled with autoclaved soil from one of two forested sites in Wisconsin, USA. Bags were buried in the site from which soil was collected. Five bags per site were collected and tested for germination and viability at six, twelve, and 22 months. To estimate the proportion that survived for approximately 17 months, we calculated the average proportion of surviving seeds (germinated or viable) between the 12 month and 22 month collection times.

To estimate survival of annual seeds that remain in the seed bank, $s_A$, we used data from the study by Redwood et al. (18). Briefly, 120 nylon mesh bags of 100 seeds each were buried in forested sites in southeastern Ohio. At twelve time points over the following two years, ten seed bags were harvested and tested for germination and viability. We extracted data from the figure of number of seeds surviving (sum of lab-germinated and non-germinated, but viable) vs. time (Fig. 2A) to estimate the number of non-germinating seeds that survived in the bags for 15 months in the absence of disease.

We did not find a significant effect of fungicide on biomass in our experiment, suggesting that the values for biomass virulence, $v_i$, are low. We therefore set these values to $1e^{-4}$, or approximately 1.6% of all infected biomass over the growing season, for all plant groups. We did not have empirical data to support estimates of transmission from litter to plants, $\beta_{jC}$, or loss of conidia from litter, *a*. We set $\beta_{jC}$ to $1\times10^{-7}$ and *a* to 0.95 so that transmission among plants would dominate disease dynamics rather than transmission from litter to live plants.

*Sensitivity analysis*

We evaluated the sensitivity of our results to parameter values by varying select parameters over a meaningful range of values and calculating invader impact (equilibrium competitor biomass pre-invasion in the default simulation, Fig. 5, divided by competitor biomass at end of simulation). We only varied parameters after disease was introduced to the system because we were most interested in the effects of disease emergence on invader impact. Parameter ranges were chosen to represent biologically valid values (based on our field estimates), but extended slightly outside of field-estimated values to characterize invader impact at the extremes.

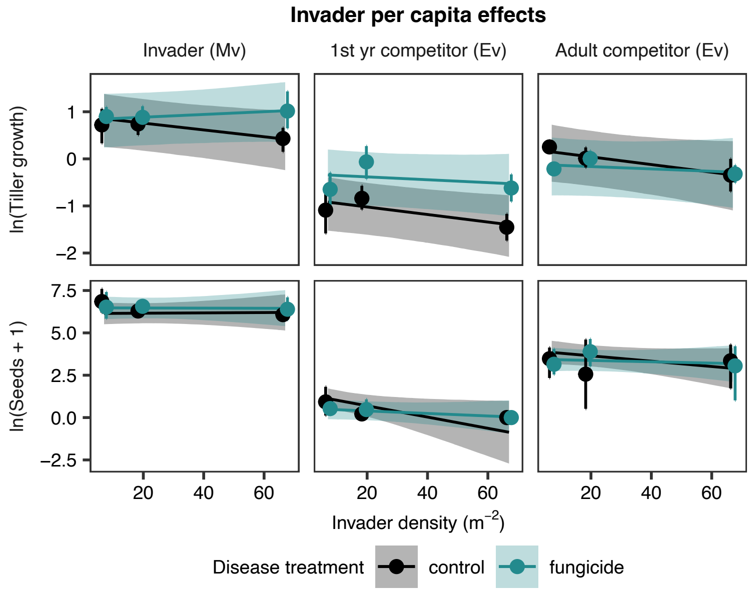


**Figure S1.** The effect of foliar fungal disease suppression on invader (Mv) per capita effects on itself and the competitor (Ev) in the first repetition of the experiment. Per capita effects were measured as the change in tiller number (ratio of numbers recorded over four-week interval) and seed production (both natural log-transformed) across the invader density gradient. To suppress foliar fungal disease, plants were sprayed with fungicide, as indicated by different colors. Points and error bars represent mean ± 95% bootstrapped confidence intervals of raw data. Lines and shading represent model-estimated mean ± 95% credible intervals.

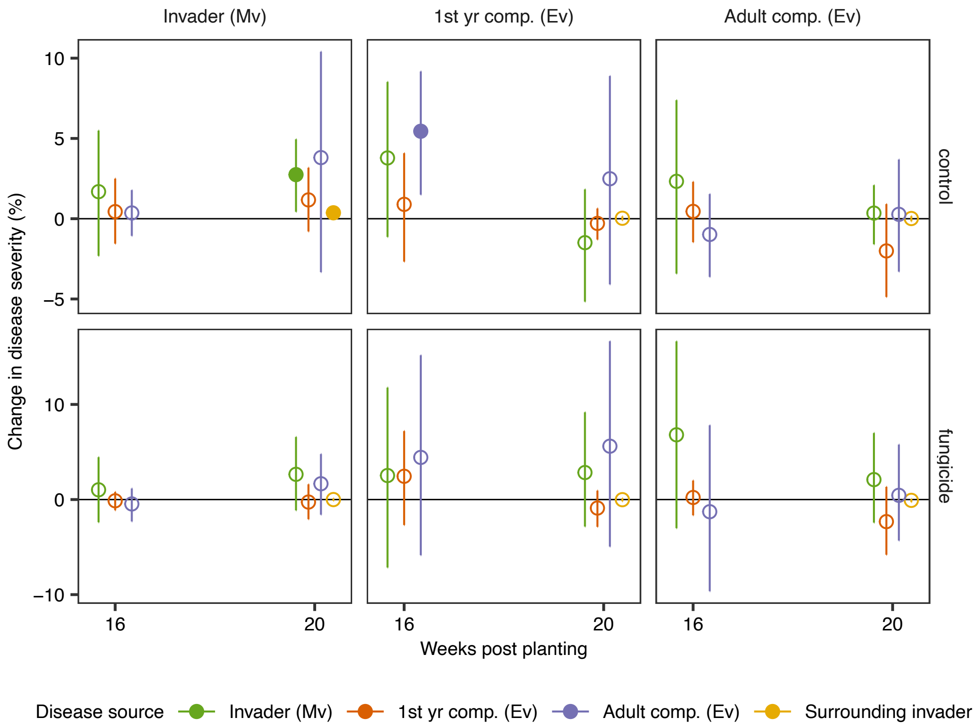


**Figure S2.** Disease transmission throughout the growing season of the first experiment repetition, measured as change in disease severity (percentage of leaf area covered by lesions) on the invader, adult competitor, and first-year competitor (panel columns) with one unit increase in disease source four weeks prior. Disease sources included the density-scaled disease severity of invader, first-year competitor, and adult competitor plants, as well as disease severity of invader plants surrounding plots. One unit of density-scaled disease severity is one plant with 100% disease severity or 100 plants with 1% disease severity and one unit of surrounding invader (which occurred at high density) is an average disease severity of 1%. Model-estimated mean and 95% credible intervals are shown. Filled points have 95% credible intervals that omit zero.

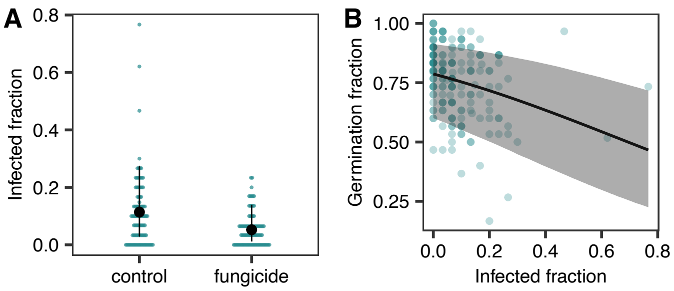


**Figure S3.** Effect of seed infection on invader seed germination. (A) Fungicide reduced the proportion of invader seeds with dark hyphae fungal infections (Table S17). (B) The proportion of invader seeds that germinated decreased with the proportion of seeds that were infected with dark hyphae fungi (Table S18). Large points and error bars (A) and lines and shading (B) represent model fits (mean ± 95% CI) and points represent raw data. Small points are binned by 0.01 to show distribution in A.

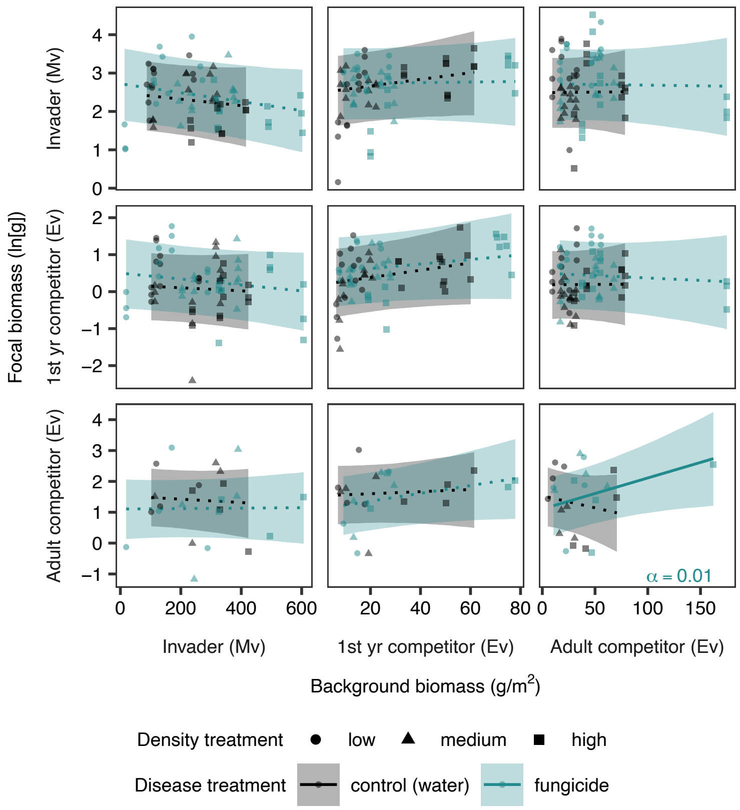


**Figure S4.** Interactions (competition and facilitation) among invader and competitor plants during the second repetition of the experiment, as estimated by log-transformed focal plant biomass over gradients of invader, adult competitor, and first-year competitor background biomass (achieved through density manipulations). Raw data are represented by points and model fits are represented by lines and shading (mean ± 95% credible intervals). Statistically significant interaction coefficients are indicated by solid lines and text values reporting the slope (α). When the largest adult competitor background biomass value is removed from the bottom right panel, the slope is still positive (α = 0.013), but not statistically different from zero.

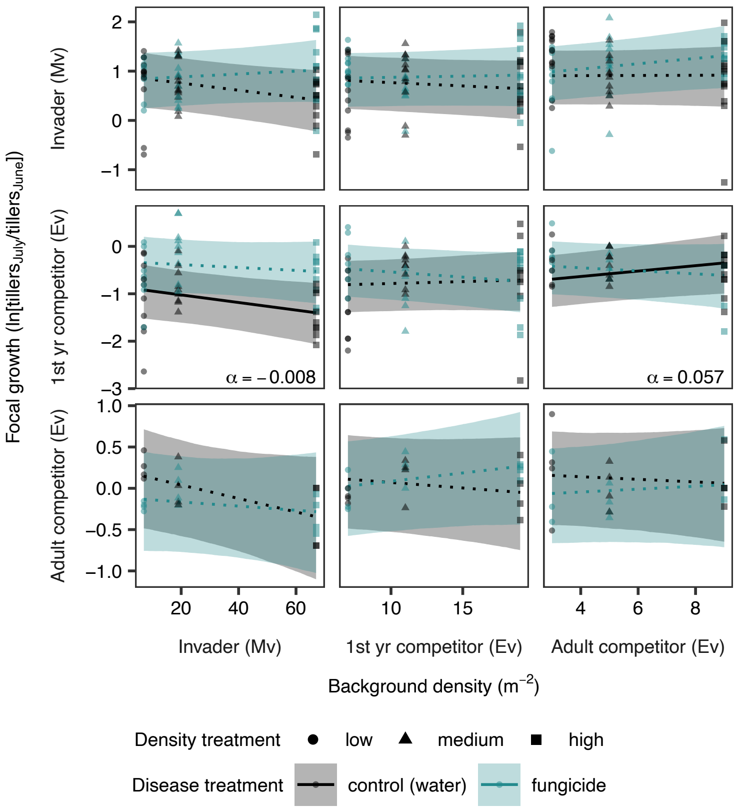


**Figure S5.** Interactions (competition and facilitation) among invader and competitor plants during the first repetition of the experiment, as estimated by focal plant growth—log transformed ratio of tillers in July over tillers in June—over gradients of invader, adult competitor, and first-year competitor density. Raw data are represented by points and model fits are represented by lines and shading (mean ± 95% credible intervals). Statistically significant interaction coefficients are indicated by solid lines and text values reporting the slope (α).

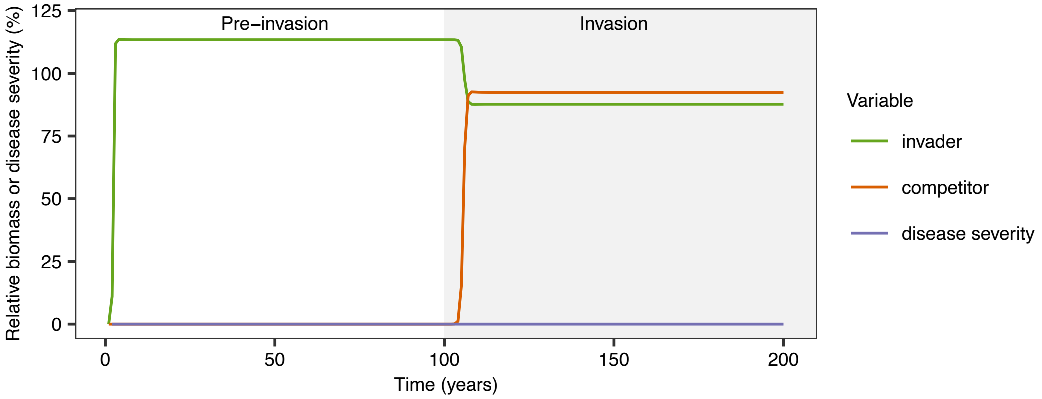


**Figure S6.** A model-simulated time-series with parameter values based on field measurements in which the competitor invades an established invader population. Relative biomass is biomass divided by the equilibrium biomass of the competitor pre-invasion (Fig. 5).

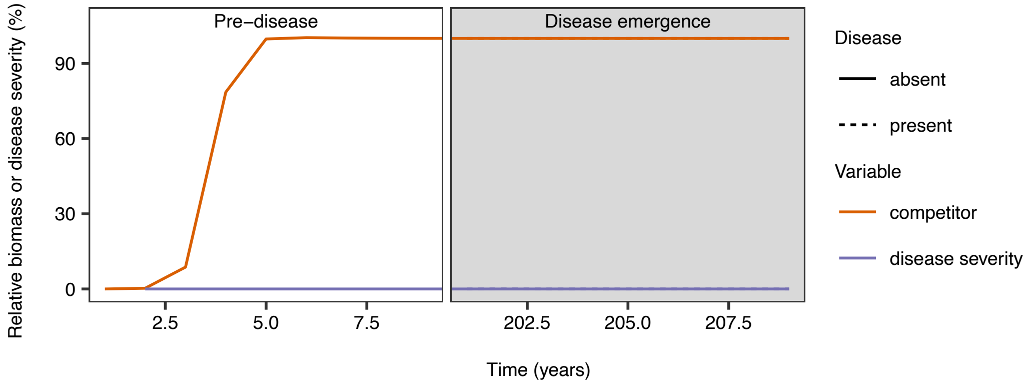


**Figure S7.** A model-simulated time-series with parameter values based on field measurements in which disease emerges in an established competitor population (years between 10 and 200 are omitted for visualization). Relative biomass is biomass divided by the equilibrium biomass of the competitor pre-invasion (Fig. 5).

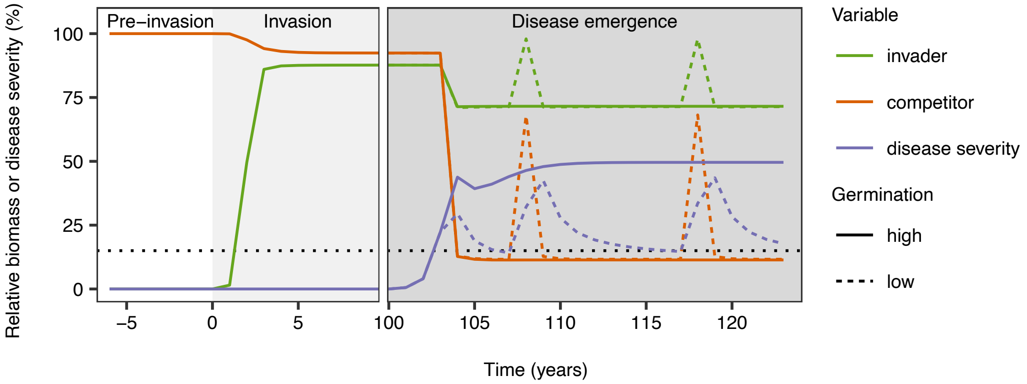


**Figure S8.** Simulations of population dynamics of the invader and competitor when invader germination was reduced due to seed infections (-44%, low germination). Simulation with high germination is the same as that shown in Fig. 5. The threshold for community-wide disease severity is indicated with a horizontal dotted black line. Parameters derived from the density experiment were set to values estimated from control plots (including reduced invader seed germination) when community-wide disease severity exceeded this threshold and fungicide-treated plots otherwise.

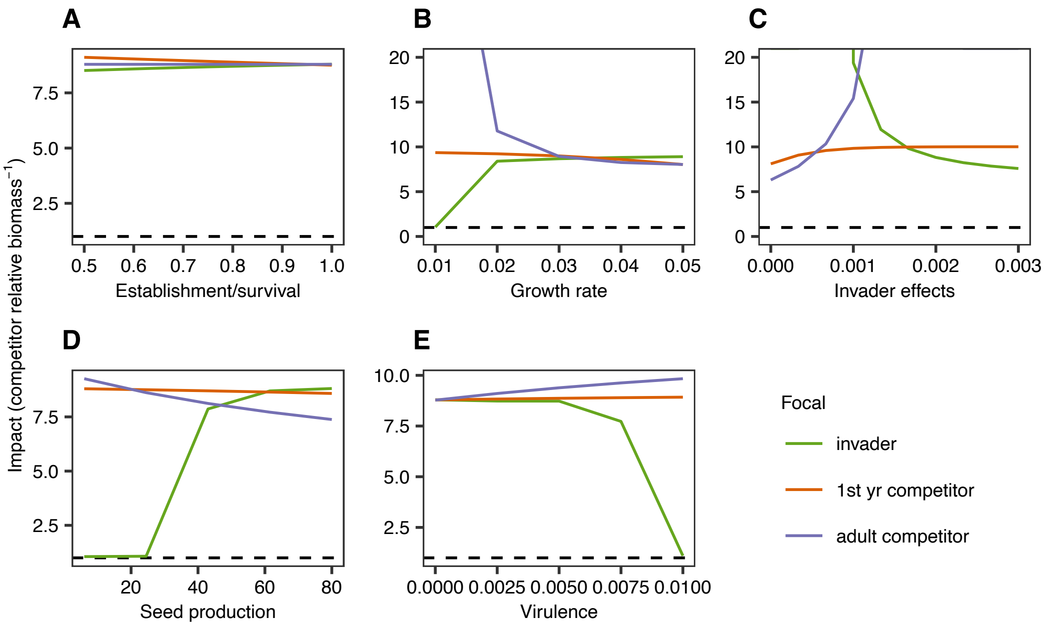


**Figure S9.** Sensitivity of impact (competitor's pre-invasion equilibrium biomass from Fig. 5 divided by competitor's final biomass) to variation in parameter values, where a value of one (dashed line) indicates no impact. (A) Greater maximum establishment fraction of invader seedlings ($e_A$) slightly increased impact while greater maximum establishment fraction of first-year competitor seedlings ($e_P$) slightly decreased impact, and adult competitor survival fraction ($l_P$) did not affect impact. (B) Greater invader growth rate ($r_A$) increased impact, greater first-year competitor growth rate ($r_F$) slightly decreased impact, and greater adult competitor growth rate ($r_P$) decreased impact. (C) Greater invader effects on invaders ($\alpha_{AA}$) decreased impact, greater invader effects on first-year competitors ($\alpha_{FA}$) slightly increased impact, and greater invader effects on adult competitors ($\alpha_{PA}$) increased impact. (D) Greater invader seed production per gram biomass ($c_A$) increased impact, greater first-year competitor seed production per gram biomass ($c_F$) did not affect impact, and greater adult competitor seed production per gram biomass ($c_P$) slightly decreased impact. (E) Greater invader virulence ($v_A$) decreased impact, greater first-year competitor virulence ($v_F$) did not affect impact, and greater adult competitor virulence ($v_P$) increased impact. Y-axes are cut-off at 20 for visualization.

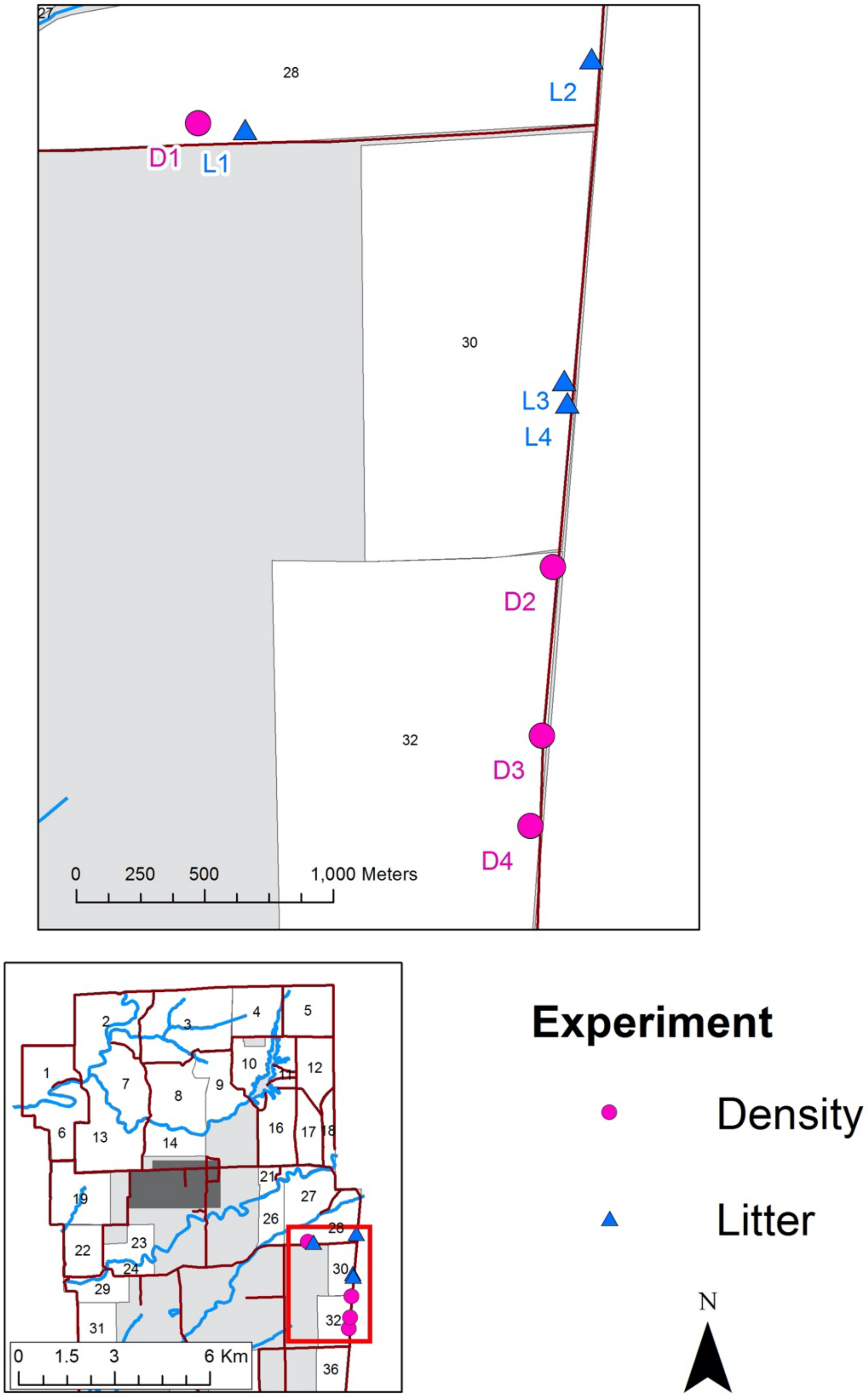


**Figure S10.** Locations of experimental sites in Big Oaks National Wildlife Refuge in Madison, IN, USA. Sites either had density manipulations (experiment described in the main text) and begin with “D” or litter manipulations (first litter experiment) and begin with “L”. Numbers and shading in maps correspond with management areas of BONWR (U.S. Fish and Wildlife Service).

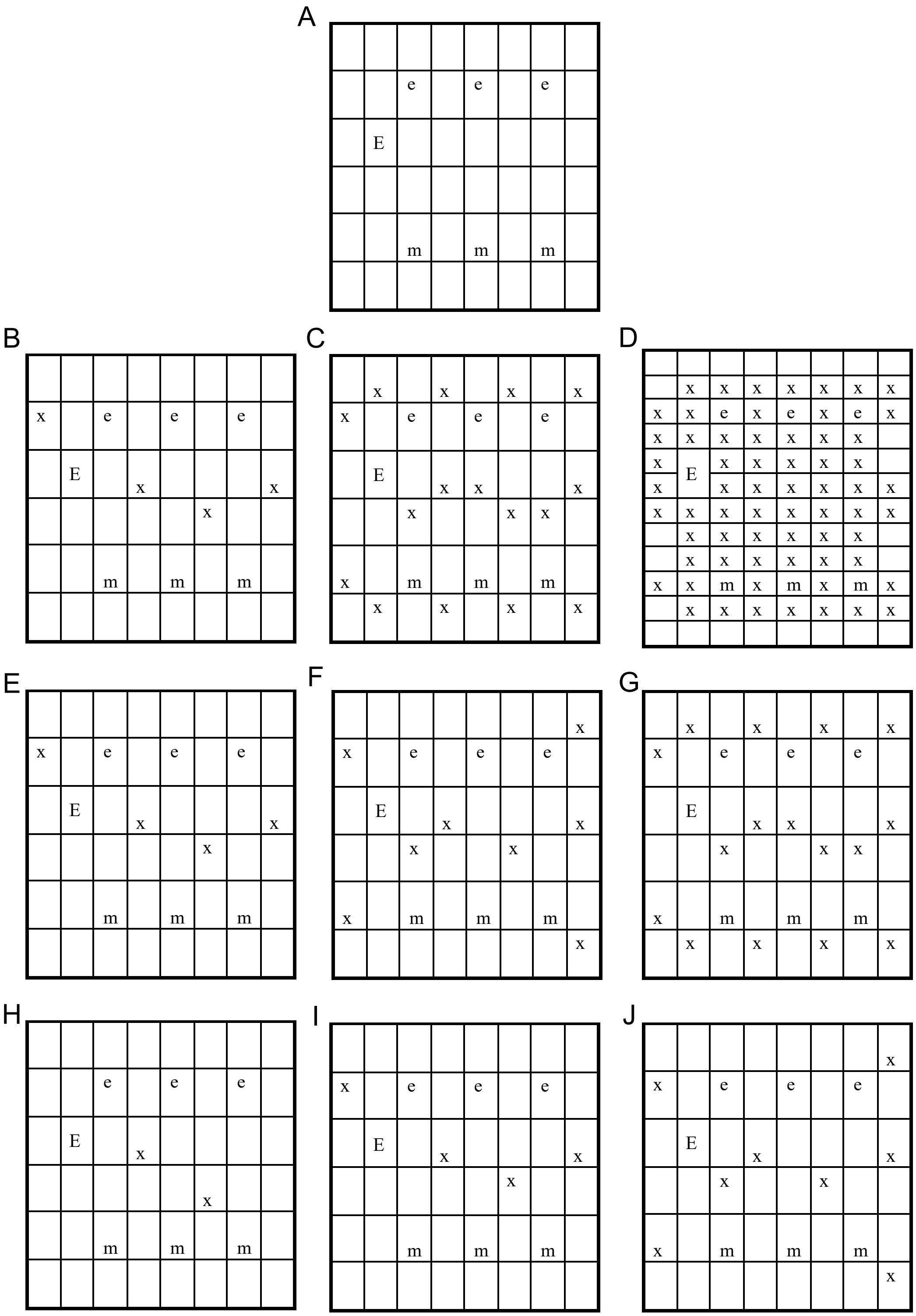


**Figure S11.** Plants positions in center 1-$m^2$ of plots with no background plants (A), low (B), medium (C), or high (D) background *M. vimineum* density, low (E), medium (F), or high (G) background *E. virginicus* first-year density, or low (H), medium (I), or high (J) background *E. virginicus* adult density. x = background plant, m = focal *M. vimineum*, e = focal *E. virginicus* first-year, E = focal *E. virginicus* adult.

**Table S1.** Model summary of plot-scale invader biomass across the invader density gradient (*N* = 32). Normal mixed-effects non-linear regression: biomass = (density × $b_0$)/(1 + α × density), where $b_0$ is maximum biomass (in grams) and α is the per capita effect of the invader on biomass.

| Variable | Estimate | Est. Error | l-95% CI | u-95% CI |
|---|---|---|---|---|
| $b_0$ control | 15.90 | 3.95 | 9.00 | 24.4 |
| $b_0$ fungicide | 14.99 | 3.89 | 8.23 | 23.32 |
| α control | 0.09 | 0.03 | 0.04 | 0.16 |
| α fungicide | 0.06 | 0.02 | 0.02 | 0.11 |
| $b_0$ random intercept: site | 26.69 | 21.90 | 2.92 | 82.68 |
| sigma | 81.37 | 11.88 | 62.30 | 108.69 |

Notes: Prior distributions were: $b_0$ – gamma (location = 14, scale = 1); α – exponential (rate = 0.5); random intercept and sigma – half Student's *t* (location = 0, df = 3, scale = 172.1)

**Table S2.** Model summary of plot-scale invader seeds across the invader density gradient (*N* = 32). Normal mixed-effects non-linear regression: seeds = (density $\times$ $s_0$)/(1 + α $\times$ density), where $s_0$ is maximum seed production and α is the per capita effect of the invader on seeds.

| Variable | Estimate | Est. Error | l-95% CI | u-95% CI |
|---|---|---|---|---|
| $s_0$ control | 1089.57 | 99.08 | 894.79 | 1283.67 |
| $s_0$ fungicide | 1049.97 | 98.58 | 854.7 | 1242.11 |
| α control | 0.06 | 0.03 | 0.03 | 0.13 |
| α fungicide | 0.07 | 0.03 | 0.03 | 0.14 |
| $s_0$ random intercept: site | 1059.30 | 1041.43 | 99.01 | 3728.89 |
| sigma | 4577.55 | 674.63 | 3484.76 | 6132.84 |

Notes: Prior distributions were: $b_0$ – normal (location = 1050, scale = 100); α – exponential (rate = 0.5); random intercept and sigma – half Student's *t* (location = 0, df = 3, scale = 10189.7)

**Table S3.** Competition coefficients based on log-transformed plant biomass across the invader and competitor density gradients (*N* = 500). See Table S4 for model details.

| Focal | Background | Treatment | Estimate | Est. Error | l-95% CI | u-95% CI |
|---|---|---|---|---|---|---|
| invader | invader | control | -0.01 | 0.01 | -0.02 | 0.00 |
| invader | invader | fungicide | -0.01 | 0.01 | -0.03 | 0.00 |
| invader | first-year comp. | control | 0.01 | 0.02 | -0.03 | 0.04 |
| invader | first-year comp. | fungicide | -0.02 | 0.02 | -0.06 | 0.01 |
| invader | adult comp. | control | -0.01 | 0.04 | -0.09 | 0.06 |
| invader | adult comp. | fungicide | -0.03 | 0.04 | -0.10 | 0.05 |
| first-year comp. | invader | control | -0.01 | 0.01 | -0.02 | 0.00 |
| first-year comp. | invader | fungicide | -0.01 | 0.01 | -0.02 | 0.00 |
| first-year comp. | first-year comp. | control | 0.01 | 0.02 | -0.03 | 0.04 |
| first-year comp. | first-year comp. | fungicide | -0.01 | 0.02 | -0.05 | 0.02 |
| first-year comp. | adult comp. | control | -0.03 | 0.04 | -0.10 | 0.04 |
| first-year comp. | adult comp. | fungicide | -0.04 | 0.04 | -0.12 | 0.03 |
| adult comp. | invader | control | -0.01 | 0.01 | -0.02 | 0.00 |
| adult comp. | invader | fungicide | 0.00 | 0.01 | -0.02 | 0.01 |
| adult comp. | first-year comp. | control | -0.02 | 0.02 | -0.06 | 0.03 |
| adult comp. | first-year comp. | fungicide | 0.02 | 0.02 | -0.02 | 0.07 |
| adult comp. | adult comp. | control | -0.12 | 0.05 | -0.21 | -0.02 |
| adult comp. | adult comp. | fungicide | 0.07 | 0.05 | -0.03 | 0.17 |

**Table S4.** Model summary of log-transformed plant biomass across the invader and competitor density gradients ($N$ = 500). Normal mixed-effects linear regression: ln(biomass) ~ focal * fungicide * (density + density:background), where focal and background (bckgrd) each take on the values of invader, first-year (1st yr) competitor, and adult competitor and fungicide was a binary variable.

| Variable | Est. | Est. Error | l-95% CI | u-95% CI |
|---|---|---|---|---|
| intercept | 2.62 | 0.19 | 2.25 | 2.99 |
| adult comp. focal | -0.78 | 0.22 | -1.21 | -0.35 |
| 1st yr comp. focal | -2.26 | 0.16 | -2.57 | -1.94 |
| fungicide | 0.30 | 0.26 | -0.21 | 0.81 |
| density | -0.01 | 0.01 | -0.02 | 0.00 |
| adult comp. focal:fungicide | -0.94 | 0.31 | -1.56 | -0.35 |
| 1st yr comp. focal:fungicide | 0.06 | 0.22 | -0.38 | 0.5 |
| density:adult comp. bckgrd | 0.00 | 0.04 | -0.07 | 0.07 |
| density: 1st yr comp. bckgrd | 0.02 | 0.02 | -0.01 | 0.05 |
| adult comp. focal:density | 0.00 | 0.01 | -0.01 | 0.01 |
| 1st yr comp. focal:density | 0.00 | 0.00 | -0.01 | 0.01 |
| fungicide:density | 0.00 | 0.01 | -0.02 | 0.01 |
| adult comp. focal:density:adult comp. bckgrd | -0.10 | 0.04 | -0.19 | -0.02 |
| 1st yr comp. focal:density:adult comp. bckgrd | -0.02 | 0.03 | -0.08 | 0.04 |
| adult comp. focal:density:1st yr comp. bckgrd | -0.02 | 0.02 | -0.06 | 0.01 |
| 1st yr comp. focal:density:1st yr comp. bckgrd | -0.01 | 0.01 | -0.03 | 0.02 |
| fungicide:density:adult comp. bckgrd | -0.01 | 0.05 | -0.11 | 0.09 |
| fungicide:density:1st yr comp. bckgrd | -0.03 | 0.02 | -0.07 | 0.01 |
| adult comp. focal:fungicide:density | 0.01 | 0.01 | -0.01 | 0.03 |
| 1st yr comp. focal:fungicide:density | 0.00 | 0.01 | -0.01 | 0.01 |
| adult comp. focal:fungicide:density:adult comp. bckgrd | 0.19 | 0.06 | 0.07 | 0.31 |
| 1st yr comp. focal:fungicide:density:adult comp. bckgrd | 0.00 | 0.04 | -0.08 | 0.09 |
| adult comp. focal:fungicide:density:1st yr comp. bckgrd | 0.06 | 0.03 | 0.01 | 0.11 |
| 1st yr comp. focal:fungicide:density:1st yr comp. bckgrd | 0.02 | 0.02 | -0.02 | 0.05 |
| random intercept: plot | 0.44 | 0.05 | 0.35 | 0.55 |
| sigma | 0.59 | 0.02 | 0.55 | 0.63 |

Notes: Prior distributions were: intercept – normal (location = 3, scale = 1); coefficients – normal (location = 0, scale = 1); random intercept and sigma – half Student's *t* (location = 0, df = 3, scale = 2.5)

**Table S5.** Competition coefficients based on log-transformed plant seed production across the invader and competitor density gradients ($N$ = 504). See Table S6 for model details.

| Focal | Background | Treatment | Estimate | Est. Error | l-95% CI | u-95% CI |
|---|---|---|---|---|---|---|
| invader | invader | control | -0.02 | 0.01 | -0.04 | 0.00 |
| invader | invader | fungicide | -0.01 | 0.01 | -0.03 | 0.00 |
| invader | first-year comp. | control | 0.02 | 0.03 | -0.04 | 0.08 |
| invader | first-year comp. | fungicide | 0.00 | 0.03 | -0.05 | 0.06 |
| invader | adult comp. | control | -0.05 | 0.06 | -0.17 | 0.06 |
| invader | adult comp. | fungicide | 0.02 | 0.06 | -0.10 | 0.14 |
| first-year comp. | invader | control | -0.01 | 0.01 | -0.03 | 0.00 |
| first-year comp. | invader | fungicide | -0.01 | 0.01 | -0.03 | 0.00 |
| first-year comp. | first-year comp. | control | -0.02 | 0.03 | -0.07 | 0.04 |
| first-year comp. | first-year comp. | fungicide | -0.03 | 0.03 | -0.09 | 0.03 |
| first-year comp. | adult comp. | control | -0.04 | 0.06 | -0.16 | 0.09 |
| first-year comp. | adult comp. | fungicide | -0.02 | 0.06 | -0.14 | 0.11 |
| adult comp. | invader | control | -0.02 | 0.01 | -0.04 | 0.01 |
| adult comp. | invader | fungicide | 0.00 | 0.01 | -0.02 | 0.03 |
| adult comp. | first-year comp. | control | -0.03 | 0.04 | -0.11 | 0.05 |
| adult comp. | first-year comp. | fungicide | 0.04 | 0.04 | -0.05 | 0.12 |
| adult comp. | adult comp. | control | -0.09 | 0.09 | -0.26 | 0.08 |
| adult comp. | adult comp. | fungicide | 0.15 | 0.09 | -0.03 | 0.33 |

**Table S6.** Model summary of log-transformed plant seed production across the invader and competitor density gradients ($N$ = 504). Normal mixed-effects linear regression: ln(seeds + 1) ~ focal * fungicide * (density + density:background), where focal and background (bckgrd) each take on the values of invader, first-year (1st yr) competitor, and adult competitor and fungicide was a binary variable.

| Variable | Est. | Est. Error | l-95% CI | u-95% CI |
|---|---|---|---|---|
| intercept | 6.62 | 0.29 | 6.05 | 7.20 |
| 1st yr comp. focal | -4.43 | 0.30 | -5.02 | -3.83 |
| adult comp. focal | -2.21 | 0.40 | -2.99 | -1.43 |
| fungicide | 0.02 | 0.39 | -0.74 | 0.77 |
| density | -0.02 | 0.01 | -0.04 | 0.00 |
| 1st yr comp. focal:fungicide | -0.13 | 0.41 | -0.93 | 0.67 |
| adult comp. focal:fungicide | -1.14 | 0.53 | -2.16 | -0.08 |
| density:1st yr comp. bckgrd | 0.04 | 0.03 | -0.01 | 0.09 |
| density:adult comp. bckgrd | -0.04 | 0.06 | -0.15 | 0.07 |
| 1st yr comp. focal:density | 0.00 | 0.01 | -0.01 | 0.02 |
| adult comp. focal:density | 0.00 | 0.01 | -0.02 | 0.03 |
| fungicide:density | 0.00 | 0.01 | -0.02 | 0.03 |
| 1st yr comp. focal:density:1st yr comp. bckgrd | -0.04 | 0.03 | -0.10 | 0.01 |
| adult comp. focal:density:1st yr comp. bckgrd | -0.05 | 0.04 | -0.12 | 0.03 |
| 1st yr comp. focal:density:adult comp. bckgrd | 0.01 | 0.06 | -0.11 | 0.13 |
| adult comp. focal:density:adult comp. bckgrd | -0.04 | 0.08 | -0.20 | 0.13 |
| fungicide:density:1st yr comp. bckgrd | -0.02 | 0.04 | -0.09 | 0.05 |
| fungicide:density:adult comp. bckgrd | 0.07 | 0.08 | -0.09 | 0.22 |
| 1st yr comp. focal:fungicide:density | 0.00 | 0.01 | -0.03 | 0.02 |
| adult comp. focal:fungicide:density | 0.02 | 0.02 | -0.02 | 0.05 |
| 1st yr comp. focal:fungicide:density:1st yr comp. bckgrd | 0.01 | 0.04 | -0.07 | 0.08 |
| adult comp. focal:fungicide:density:1st yr comp. bckgrd | 0.06 | 0.05 | -0.04 | 0.16 |
| 1st yr comp. focal:fungicide:density:adult comp. bckgrd | -0.05 | 0.08 | -0.21 | 0.12 |
| adult comp. focal:fungicide:density:adult comp. bckgrd | 0.16 | 0.12 | -0.07 | 0.38 |
| random intercept: plot | 0.60 | 0.09 | 0.44 | 0.78 |
| sigma | 1.21 | 0.04 | 1.13 | 1.30 |

Notes: Prior distributions were: intercept – normal (location = 7, scale = 1); coefficients – normal (location = 0, scale = 1); random intercept and sigma – half Student's *t* (location = 0, df = 3, scale = 3.1)

**Table S7.** The effect of fungicide (percent change relative to control) on the per capita effects of the invader in the second year of the experiment, measured as the change in tillers over a four-week interval ($N$ = 416) and the number of seeds produced ($N$ = 244). Normal mixed-effects linear regressions: ln(tiller growth) or ln(seeds + 1) ~ focal * fungicide * (density + density:background), where focal and background each take on the values of invader, first-year competitor, and adult competitor and fungicide was a binary variable.

| Response | Focal | Estimate (%) | l-95% CI | u-95% CI |
|---|---|---|---|---|
| tiller growth | adult comp. | -74 | -485 | 360 |
| tiller growth | first-year comp. | -63 | -246 | 246 |
| tiller growth | invader | -137 | -463 | 56 |
| seeds | adult comp. | -79 | -549 | 368 |
| seeds | first-year comp. | -79 | -168 | 62 |
| seeds | invader | -90 | -1298 | 1191 |

Notes: Prior distributions were (in order of growth/seed): intercept – normal (location = 0.75, scale = 1)/(location = 7, scale = 1); coefficients – normal (location = 0, scale = 1); random intercept and sigma – half Student's *t* (location = 0, df = 3, scale = 2.5)/(location = 0, df = 3, scale = 4.4)

**Table S8.** Model summary of plot-scale invader emergence across the invader litter gradient ($N$ = 28). Binomial (logit-link) mixed-effects generalized linear regression: proportion emerged ~ litter weight * autoclaved, where autoclaved was a binary variable and litter weight was in units of $g/m^2$.

| Variable | Estimate | Est. Error | l-95% CI | u-95% CI |
|---|---|---|---|---|
| intercept | 0.65 | 0.32 | -0.01 | 1.30 |
| litter | -39.41 | 4.05 | -47.37 | -31.57 |
| autoclaved | -0.03 | 0.06 | -0.15 | 0.10 |
| litter:autoclaved | -5.21 | 5.02 | -15.09 | 4.45 |
| random intercept: site | 0.55 | 0.41 | 0.19 | 1.65 |

Notes: Prior distributions were: intercept and coefficients – normal (location = 0, scale = 10), random intercept – half Student's *t* (location = 0, df = 3, scale = 2.5)

**Table S9.** Model summary of plot-scale competitor emergence across the invader litter gradient ($N$ = 28). Normal non-linear regression: proportion emerged = $e$ / (1 + γ × litter), where $e$ is the maximum proportion emerged and γ is the per-gram effect of litter on emergence.

| Variable | Estimate | Est. Error | l-95% CI | u-95% CI |
|---|---|---|---|---|
| $e$ | 0.03 | 0.01 | 0.00 | 0.06 |
| γ | 12.64 | 11.37 | 0.41 | 41.73 |
| sigma | 0.03 | 0.01 | 0.02 | 0.05 |

Notes: Prior distributions were: $e$ – normal (location = 0.04, scale = 1), γ – exponential (rate = 0.1), sigma – half Student's $t$ (location = 0, df = 3, scale = 2.5)

**Table S10.** Model summaries of disease severity at 8 ($N$ = 538), 12 ($N$ = 514), and 16 ($N$ = 456) weeks post planting. Beta (logit-link) mixed-effects generalized linear regression: disease severity ~ focal * fungicide * (src invader + src 1$^{st}$ yr competitor + src adult competitor + src surr), where focal took on the values of invader, first-year competitor (1$^{st}$ yr comp), and adult competitor (adult comp). Disease sources (src) were measured 4 weeks prior to the response variable as density-weighted disease severity of the invader, first-year competitors, and adult competitors in the plot as well as disease severity of invader plants surrounding (surr) the of plot.

| | 8 weeks post planting | | | 12 weeks post planting | | | 16 weeks post planting | | |
|---|---|---|---|---|---|---|---|---|---|
| Variable | Est. | l-95% | u-95% | Est. | l-95% | u-95% | Est. | l-95% | u-95% |
| Intercept | -3.82 | -4.17 | -3.49 | -3.12 | -3.44 | -2.81 | -1.32 | -1.65 | -1.01 |
| 1st yr comp | 1.08 | 0.72 | 1.44 | 1.73 | 1.38 | 2.08 | 1.28 | 0.86 | 1.70 |
| adult comp | 1.33 | 0.85 | 1.78 | 1.88 | 1.41 | 2.34 | 1.58 | 0.89 | 2.27 |
| fung | -0.02 | -0.39 | 0.36 | -0.41 | -0.82 | -0.01 | -0.62 | -1.04 | -0.19 |
| src invader | -0.04 | -0.34 | 0.20 | -0.31 | -2.22 | 1.58 | 0.06 | -0.06 | 0.19 |
| src 1st yr | -0.03 | -0.23 | 0.16 | 0.45 | 0.21 | 0.68 | -0.08 | -0.19 | 0.02 |
| src adult | -0.06 | -0.59 | 0.45 | 0.61 | 0.04 | 1.14 | -0.32 | -0.76 | 0.11 |
| src surr | 0.23 | -0.36 | 0.79 | 0.59 | -0.12 | 1.28 | 0.32 | 0.19 | 0.46 |
| 1st yr comp:fung | -0.16 | -0.62 | 0.29 | -0.46 | -0.95 | 0.04 | 0.78 | 0.24 | 1.31 |
| adult comp:fung | 0.22 | -0.35 | 0.80 | 0.11 | -0.51 | 0.74 | 0.55 | -0.29 | 1.38 |
| 1st yr comp:src invader | -0.09 | -0.44 | 0.25 | -0.22 | -2.14 | 1.68 | 0.09 | -0.07 | 0.26 |
| adult comp:src invader | -0.04 | -0.67 | 0.44 | -0.35 | -2.25 | 1.56 | 0.23 | -0.06 | 0.55 |
| 1st yr comp:src 1st yr | 0.01 | -0.22 | 0.24 | -0.58 | -0.86 | -0.29 | 0.13 | -0.03 | 0.31 |
| adult comp:src 1st yr | 0.08 | -0.28 | 0.42 | -0.53 | -0.95 | -0.13 | -0.20 | -0.73 | 0.32 |
| 1st yr comp:src adult | 0.52 | -0.06 | 1.10 | -1.05 | -1.72 | -0.37 | 0.18 | -0.41 | 0.76 |
| adult comp:src adult | -0.54 | -1.47 | 0.34 | -0.72 | -1.77 | 0.29 | 0.32 | -0.84 | 1.46 |
| 1st yr comp:src surr | 0.9 | 0.25 | 1.57 | -0.78 | -1.6 | 0.05 | -0.41 | -0.6 | -0.23 |
| adult comp:src surr | 0.61 | -0.32 | 1.49 | -0.92 | -1.95 | 0.10 | -0.50 | -0.86 | -0.20 |
| fung:src invader | 0.15 | -1.59 | 1.84 | 0.04 | -1.93 | 1.98 | 0.09 | -1.50 | 1.66 |
| fung:src 1st yr | -0.01 | -0.30 | 0.28 | -0.42 | -0.81 | -0.04 | 0.28 | 0.05 | 0.49 |
| fung:src adult | 0.00 | -0.56 | 0.58 | -0.22 | -0.97 | 0.54 | 0.36 | -0.12 | 0.85 |
| fung:src surr | -0.24 | -0.85 | 0.38 | 0.26 | -0.64 | 1.15 | -0.10 | -0.30 | 0.09 |
| 1st yr comp:fung:src invader | 0.16 | -1.71 | 1.99 | 0.01 | -1.97 | 1.98 | 0.13 | -1.58 | 1.85 |
| adult comp:fung:src invader | 0.12 | -1.76 | 2.03 | 0.03 | -1.96 | 2.01 | -0.36 | -2.22 | 1.53 |
| 1st yr comp:fung:src 1st yr | 0.06 | -0.30 | 0.42 | 0.79 | 0.33 | 1.24 | -0.18 | -0.47 | 0.12 |
| adult comp:fung:src 1st yr | -0.12 | -0.58 | 0.35 | 0.39 | -0.32 | 1.04 | -0.02 | -0.64 | 0.62 |
| 1st yr comp:fung:src adult | -0.19 | -0.84 | 0.45 | 1.48 | 0.54 | 2.39 | -0.02 | -0.70 | 0.68 |
| adult comp:fung:src adult | 0.22 | -0.75 | 1.20 | 0.36 | -0.93 | 1.60 | -0.37 | -1.71 | 1.02 |
| 1st yr comp:fung:src surr | -0.64 | -1.36 | 0.07 | 0.23 | -1.00 | 1.43 | 0.13 | -0.13 | 0.38 |
| adult comp:fung:src surr | -0.34 | -1.28 | 0.63 | 1.02 | -0.45 | 2.48 | 0.35 | -0.05 | 0.78 |
| random intercept: plot | 0.23 | 0.03 | 0.39 | 0.27 | 0.09 | 0.41 | 0.26 | 0.08 | 0.40 |
| phi | 12.37 | 10.16 | 14.87 | 6.94 | 5.80 | 8.14 | 3.87 | 3.35 | 4.42 |

Notes: Prior distributions were: intercept and coefficients – normal (location = 0, scale = 1), phi – gamma (location = 0.01, scale = 0.01), random intercept – half Student's *t* (location = 0, df = 3, scale = 2.5)

**Table S11.** Model summaries of disease severity at 16 (*N* = 349) and 20 (*N* = 306) weeks post planting in the first repetition of the experiment. Beta (logit-link) mixed-effects generalized linear regression: disease severity ~ focal * fungicide * (src invader + src 1$^{st}$ yr competitor + src adult competitor + src surr), where focal took on the values of invader, first-year competitor (1$^{st}$ yr comp), and adult competitors (adult comp). Disease sources (src) were measured 8 (for 16 weeks) or 4 (for 20 weeks) weeks prior to the response variable as density-weighted disease severity of the invader, first-year competitors, and adult competitors in the plot as well as percentage of biomass with lesions of invader plants surrounding (surr) the outside of plots (only measured 8 weeks post planting).

| | 16 weeks post planting | | | | 20 weeks post planting | | | |
|---|---|---|---|---|---|---|---|---|
| Covariate | Est. | Error | l-95% | u-95% | Est. | Error | l-95% | u-95% |
| Intercept | -2.57 | 0.15 | -2.87 | -2.27 | -1.99 | 0.14 | -2.27 | -1.71 |
| 1st yr comp | -0.20 | 0.23 | -0.65 | 0.26 | -1.00 | 0.34 | -1.69 | -0.35 |
| adult comp | -0.24 | 0.24 | -0.71 | 0.23 | -0.75 | 0.32 | -1.39 | -0.15 |
| fung | -0.55 | 0.20 | -0.94 | -0.17 | -1.03 | 0.24 | -1.50 | -0.56 |
| src invader | 0.24 | 0.29 | -0.34 | 0.77 | 0.19 | 0.08 | 0.03 | 0.35 |
| src 1st yr | 0.06 | 0.15 | -0.24 | 0.34 | 0.08 | 0.07 | -0.06 | 0.22 |
| src adult | 0.05 | 0.10 | -0.16 | 0.24 | 0.27 | 0.24 | -0.23 | 0.73 |
| 1st yr comp:fung | 0.51 | 0.32 | -0.12 | 1.12 | 0.83 | 0.43 | -0.03 | 1.68 |
| adult comp:fung | 0.83 | 0.34 | 0.14 | 1.50 | 1.01 | 0.46 | 0.11 | 1.93 |
| 1st yr comp:src invader | 0.27 | 0.39 | -0.51 | 1.03 | -0.50 | 0.37 | -1.26 | 0.17 |
| adult comp:src invader | 0.18 | 0.50 | -0.88 | 1.10 | -0.11 | 0.20 | -0.55 | 0.23 |
| 1st yr comp:src 1st yr | 0.05 | 0.26 | -0.46 | 0.55 | -0.14 | 0.11 | -0.37 | 0.06 |
| adult comp:src 1st yr | 0.02 | 0.21 | -0.42 | 0.42 | -0.52 | 0.34 | -1.24 | 0.09 |
| 1st yr comp:src adult | 0.68 | 0.26 | 0.12 | 1.16 | 0.25 | 0.66 | -1.08 | 1.53 |
| adult comp:src adult | -0.23 | 0.26 | -0.78 | 0.23 | -0.21 | 0.40 | -1.04 | 0.51 |
| fung:src invader | 0.01 | 0.47 | -0.94 | 0.91 | 0.31 | 0.37 | -0.43 | 1.01 |
| fung:src 1st yr | -0.10 | 0.18 | -0.45 | 0.27 | -0.14 | 0.20 | -0.54 | 0.23 |
| fung:src adult | -0.17 | 0.24 | -0.67 | 0.28 | 0.06 | 0.38 | -0.69 | 0.82 |
| 1st yr comp:fung:src invader | -0.2 | 0.67 | -1.54 | 1.10 | 0.55 | 0.64 | -0.72 | 1.79 |
| adult comp:fung:src invader | 0.52 | 0.71 | -0.90 | 1.90 | 0.25 | 0.67 | -1.12 | 1.53 |
| 1st yr comp:fung:src 1st yr | 0.29 | 0.40 | -0.50 | 1.05 | 0.00 | 0.27 | -0.55 | 0.52 |
| adult comp:fung:src 1st yr | 0.05 | 0.26 | -0.45 | 0.57 | -0.34 | 0.70 | -1.74 | 1.00 |
| 1st yr comp:fung:src adult | -0.02 | 0.67 | -1.35 | 1.27 | 0.58 | 0.91 | -1.20 | 2.39 |
| adult comp:fung:src adult | 0.10 | 0.66 | -1.24 | 1.35 | 0.03 | 0.76 | -1.50 | 1.49 |
| src surr | NA | NA | NA | NA | 0.03 | 0.01 | 0.01 | 0.04 |
| 1st yr comp:src surr | NA | NA | NA | NA | -0.02 | 0.01 | -0.05 | 0.01 |
| adult comp:src surr | NA | NA | NA | NA | -0.02 | 0.02 | -0.06 | 0.01 |
| fung:src surr | NA | NA | NA | NA | -0.03 | 0.01 | -0.06 | 0.00 |
| 1st yr comp:fung:src surr | NA | NA | NA | NA | 0.02 | 0.02 | -0.03 | 0.06 |
| adult comp:fung:src surr | NA | NA | NA | NA | -0.01 | 0.03 | -0.07 | 0.06 |
| random intercept: plot | 0.15 | 0.09 | 0.01 | 0.33 | 0.29 | 0.11 | 0.04 | 0.50 |
| phi | 8.37 | 0.80 | 6.89 | 10.02 | 8.96 | 1.03 | 7.10 | 11.10 |

Notes: Prior distributions were: intercept and coefficients – normal (location = 0, scale = 1), phi – gamma (location = 0.01, scale = 0.01), random intercept – half Student's *t* (location = 0, df = 3, scale = 2.5)

**Table S12.** Coefficients for the effects of dew intensity on disease severity (%) at 12 (*N* = 258) and 16 (*N* = 222) weeks post planting in control plots during the second repetition of the experiment. Beta (logit-link) mixed-effects generalized linear regression: disease severity ~ focal * (src invader + src 1$^{st}$ yr competitor + src adult competitor + src surr + dew intensity), where focal took on the values of invader, first-year competitor, and adult competitor, fungicide was a binary variable, and disease source variables are described in Table S10. Dew intensity was centered on its mean value.

| Weeks | Focal | Estimate | l-95% CI | u-95% CI |
|---|---|---|---|---|
| 12 | invader | -0.11 | -1.43 | 1.30 |
| 12 | first-year comp. | -2.98 | -5.90 | -0.05 |
| 12 | adult comp. | -1.47 | -7.25 | 4.51 |
| 16 | invader | 0.49 | -2.95 | 3.80 |
| 16 | first-year comp. | -3.65 | -9.23 | 1.97 |
| 16 | adult comp. | 8.40 | -3.09 | 19.78 |
| | Variable | Estimate | l-95% CI | u-95% CI |
| 12 | random intercept: plot | 0.21 | 0.01 | 0.42 |
| 12 | phi | 6.19 | 4.97 | 7.57 |
| 16 | random intercept: plot | 0.17 | 0.01 | 0.38 |
| 16 | phi | 3.70 | 3.05 | 4.42 |

Notes: Prior distributions were: intercept and coefficients – normal (location = 0, scale = 1), phi – gamma (location = 0.01, scale = 0.01), random intercept – half Student's *t* (location = 0, df = 3, scale = 2.5)

**Table S13.** Coefficients for the effects of soil moisture and canopy cover on disease severity (%) at 16 ($N$ = 349) and 20 ($N$ = 306) weeks post planting in the first repetition of the experiment. Beta (logit-link) mixed-effects generalized linear regression: disease severity ~ focal * fungicide * (src invader + src 1st yr competitor + src adult competitor + src surr + canopy cover + soil moisture), where focal took on the values of invader, first-year competitor, and adult competitor, fungicide was a binary variable, and disease source values are described in Table S11. Canopy cover and soil moisture proportions were centered on their mean values.

| Weeks | Focal | Variable | Treatment | Estimate | l-95% CI | u-95% CI |
|---|---|---|---|---|---|---|
| 16 | invader | canopy cover | control | 0.00 | -0.11 | 0.12 |
| 16 | invader | canopy cover | fungicide | -0.01 | -0.08 | 0.06 |
| 16 | first-year comp. | canopy cover | control | 0.16 | -0.01 | 0.34 |
| 16 | first-year comp. | canopy cover | fungicide | -0.01 | -0.19 | 0.19 |
| 16 | adult comp. | canopy cover | control | -0.02 | -0.17 | 0.15 |
| 16 | adult comp. | canopy cover | fungicide | -0.02 | -0.20 | 0.16 |
| 16 | invader | soil moisture | control | -0.13 | -0.30 | 0.03 |
| 16 | invader | soil moisture | fungicide | -0.06 | -0.18 | 0.07 |
| 16 | first-year comp. | soil moisture | control | -0.20 | -0.50 | 0.10 |
| 16 | first-year comp. | soil moisture | fungicide | -0.44 | -0.85 | -0.02 |
| 16 | adult comp. | soil moisture | control | -0.07 | -0.31 | 0.17 |
| 16 | adult comp. | soil moisture | fungicide | 0.03 | -0.49 | 0.54 |
| 20 | invader | canopy cover | control | 0.03 | -0.22 | 0.30 |
| 20 | invader | canopy cover | fungicide | 0.04 | -0.06 | 0.14 |
| 20 | first-year comp. | canopy cover | control | -0.11 | -0.29 | 0.06 |
| 20 | first-year comp. | canopy cover | fungicide | 0.00 | -0.15 | 0.16 |
| 20 | adult comp. | canopy cover | control | -0.07 | -0.23 | 0.09 |
| 20 | adult comp. | canopy cover | fungicide | -0.01 | -0.10 | 0.08 |
| 20 | invader | soil moisture | control | -0.14 | -0.58 | 0.30 |
| 20 | invader | soil moisture | fungicide | -0.05 | -0.30 | 0.20 |
| 20 | first-year comp. | soil moisture | control | 0.09 | -0.22 | 0.40 |
| 20 | first-year comp. | soil moisture | fungicide | -0.08 | -0.41 | 0.26 |
| 20 | adult comp. | soil moisture | control | 0.08 | -0.34 | 0.49 |
| 20 | adult comp. | soil moisture | fungicide | -0.09 | -0.36 | 0.16 |

| | Variable | Estimate | l-95% CI | u-95% CI |
|---|---|---|---|---|
| 16 | random intercept: plot | 0.13 | 0.01 | 0.31 |
| 16 | phi | 8.64 | 7.14 | 10.34 |
| 20 | random intercept: plot | 0.38 | 0.15 | 0.59 |
| 20 | phi | 9.60 | 7.53 | 11.92 |

Notes: Prior distributions were: intercept and coefficients – normal (location = 0, scale = 1), phi – gamma (location = 0.01, scale = 0.01), random intercept – half Student's *t* (location = 0, df = 3, scale = 2.5)

**Table S14.** Number of sites (out of 4, unless otherwise noted in parentheses) in which *Bipolaris* infection was identified by microscopy on leaves sampled from the invader (*M. vimineum*) and competitor (*E. virginicus*).

| Experiment repetition | Weeks post planting | Treatment | Invader | Competitor |
|---|---|---|---|---|
| 1 | 8 | control | 4 | 1 |
| 1 | 8 | fungicide | 4 | 2 |
| 1 | 16 | control | 4 | 4 |
| 1 | 16 | fungicide | 4 | 3 |
| 1 | 20 | control | 3 | 1 (3) |
| 1 | 20 | fungicide | 3 | 0 (3) |
| 2 | 8 | control | 4 | NA |
| 2 | 8 | fungicide | 4 | NA |
| 2 | 12 | control | 4 | 3 (3) |
| 2 | 12 | fungicide | 4 | 3 (3) |

**Table S15.** Model summary of proportion of invader seeds that germinated (*N* = 184), measured with 1–3 samples of 30 seeds per plot. Binomial (logit-link) mixed-effects generalized linear regression: proportion germinated ~ fungicide, where fungicide was a binary variable.

| Variable | Estimate | Est. Error | l-95% CI | u-95% CI |
|---|---|---|---|---|
| intercept | 1.16 | 0.10 | 0.97 | 1.35 |
| fungicide | 0.10 | 0.14 | -0.17 | 0.37 |
| random intercept: plot | 0.53 | 0.06 | 0.42 | 0.66 |

Notes: Prior distributions were: intercept and coefficient – normal (location = 0, scale = 10), random intercept – half Student's *t* (location = 0, df = 3, scale = 2.5)

**Table S16.** Model summary of proportion of competitor seeds that germinated (*N* = 96), measured with seeds collected in both years of the experiment from adults and first-year plants growing in high density plots of invaders, first-year competitors, and adult competitors. Binomial (logit-link) mixed-effects generalized linear regression: proportion germinated ~ fungicide, where fungicide was a binary variable.

| Variable | Estimate | Est. Error | l-95% CI | u-95% CI |
|---|---|---|---|---|
| intercept | -1.52 | 1.33 | -4.15 | 1.47 |
| fungicide | -0.36 | 0.11 | -0.58 | -0.15 |
| random intercept: site | 1.10 | 0.70 | 0.41 | 2.98 |
| random intercept: year | 1.30 | 1.29 | 0.16 | 4.84 |

Notes: Prior distributions were: intercept and coefficient – normal (location = 0, scale = 10), random intercept – half Student's *t* (location = 0, df = 3, scale = 2.5)

**Table S17.** Model summary of proportion of invader seeds with dark hyphae fungal infections (*N* = 184), measured with 1–3 samples of 30 seeds per plot. Binomial (logit-link) mixed-effects generalized linear regression: proportion infected ~ fungicide, where fungicide was a binary variable.

| Variable | Estimate | Est. Error | l-95% CI | u-95% CI |
|---|---|---|---|---|
| intercept | -2.2 | 0.12 | -2.45 | -1.96 |
| fungicide | -0.87 | 0.19 | -1.24 | -0.51 |
| random intercept: plot | 0.64 | 0.09 | 0.48 | 0.83 |

Notes: Prior distributions were: intercept and coefficient – normal (location = 0, scale = 10), random intercept – half Student's *t* (location = 0, df = 3, scale = 2.5)

**Table S18.** Model summary of proportion of invader seeds that germinated ($N$ = 184), measured with 1–3 samples of 30 seeds per plot. Binomial (logit-link) mixed-effects generalized linear regression: proportion germinated ~ dark hyphae + light hyphae, where dark and light hyphae are proportions.

| Covariate | Estimate | Est. Error | l-95% CI | u-95% CI |
|---|---|---|---|---|
| intercept | 1.19 | 0.10 | 1.01 | 1.38 |
| infection with dark hyphae fungi | -1.99 | 0.36 | -2.68 | -1.28 |
| infection with light hyphae fungi | 0.79 | 0.25 | 0.30 | 1.29 |
| random intercept: plot | 0.51 | 0.06 | 0.41 | 0.63 |

Notes: Prior distributions were: intercept and coefficient – normal (location = 0, scale = 10), random intercept – half Student's *t* (location = 0, df = 3, scale = 2.5)

**Table S19.** Model summary of the proportion of seedlings that established and survived through the growing season (*N* = 560). Bernoulli (logit-link) mixed-effects generalized linear regression: proportion survived ~ fungicide * focal, where focal took on the values of invader, first-year competitor, and adult competitor and fungicide was a binary variable.

| Variable | Estimate | Est. Error | l-95% CI | u-95% CI |
|---|---|---|---|---|
| intercept | 4.84 | 0.94 | 3.31 | 6.97 |
| fungicide | 5.37 | 3.57 | 0.05 | 13.61 |
| adult competitor | -1.81 | 1.05 | -3.97 | 0.17 |
| first-year competitor | -1.89 | 0.90 | -3.86 | -0.34 |
| fungicide:adult competitor | -5.31 | 3.65 | -13.72 | 0.33 |
| fungicide:first-year competitor | -2.47 | 3.74 | -10.92 | 3.60 |
| random intercept: plot | 0.92 | 0.57 | 0.05 | 2.19 |

Notes: Prior distributions were: intercept and coefficients – normal (location = 0, scale = 10), random intercept – half Student's *t* (location = 0, df = 3, scale = 2.5)

**Table S20.** Model summary of log-transformed competitor biomass (*N* = 19) from the greenhouse experiment (Methods S2). Normal linear regression: log(biomass) ~ fungicide, where fungicide was a binary variable.

| Variable | Estimate | Est. Error | l-95% CI | u-95% CI |
|---|---|---|---|---|
| intercept | 2.58 | 0.06 | 2.46 | 2.71 |
| fungicide | 0.03 | 0.09 | -0.13 | 0.21 |
| sigma | 0.18 | 0.03 | 0.13 | 0.26 |

Notes: Prior distributions were: intercept and coefficients – normal (location = 0, scale = 10), sigma – half Student's *t* (location = 0, df = 3, scale = 2.5)

**Table S21.** Model summary of log-transformed invader biomass ($N$ = 19) from the greenhouse experiment (Methods S2). Normal linear regression: log(biomass) ~ fungicide, where fungicide was a binary variable.

| Variable | Estimate | Est. Error | l-95% CI | u-95% CI |
|---|---|---|---|---|
| intercept | 3.16 | 0.1 | 2.96 | 3.36 |
| fungicide | -0.16 | 0.14 | -0.45 | 0.12 |
| sigma | 0.30 | 0.06 | 0.22 | 0.44 |

Notes: Prior distributions were: intercept and coefficients – normal (location = 0, scale = 10), sigma – half Student's *t* (location = 0, df = 3, scale = 2.5)

**Table S22.** Model summary of invader seed heads (*N* = 19) from the greenhouse experiment (Methods S2). Negative binomial (log-link) generalized linear regression: seeds heads ~ fungicide, where fungicide was a binary variable

| Variable | Estimate | Est. Error | l-95% CI | u-95% CI |
|---|---|---|---|---|
| intercept | 3.6 | 0.09 | 3.42 | 3.78 |
| fungicide | -0.21 | 0.12 | -0.46 | 0.03 |
| shape | 35.79 | 29.83 | 9.03 | 113.49 |

Notes: Prior distributions were: intercept and coefficients – normal (location = 0, scale = 10), shape – gamma (location = 0.01, scale = 0.01)

**Table S23.** Model summary of log-transformed plant biomass across invader and competitor biomass gradients (*N* = 500). Normal mixed-effects linear regression: ln(biomass) ~ focal * fungicide * (biomass + biomass:background), where focal and background each take on the values of invader, first-year competitor, and adult competitor and fungicide was a binary variable.

| Variable | Estimate | Est. Error | l-95% CI | u-95% CI |
|---|---|---|---|---|
| intercept | 2.50 | 0.18 | 2.14 | 2.86 |
| adult comp. focal | -0.96 | 0.21 | -1.37 | -0.55 |
| first-year comp. focal | -2.30 | 0.15 | -2.60 | -2.00 |
| fungicide | 0.23 | 0.24 | -0.23 | 0.70 |
| biomass | 0.00 | 0.00 | 0.00 | 0.00 |
| adult comp. focal:fungicide | -0.66 | 0.28 | -1.21 | -0.11 |
| first-year comp. focal:fungicide | 0.08 | 0.21 | -0.31 | 0.49 |
| biomass:adult comp. background | 0.00 | 0.01 | -0.01 | 0.01 |
| biomass:first-year comp. background | 0.01 | 0.01 | 0.00 | 0.02 |
| adult comp. focal:biomass | 0.00 | 0.00 | 0.00 | 0.00 |
| first-year comp. focal:biomass | 0.00 | 0.00 | 0.00 | 0.00 |
| fungicide:biomass | 0.00 | 0.00 | 0.00 | 0.00 |
| adult comp. focal:biomass:adult comp. background | -0.01 | 0.01 | -0.02 | 0.01 |
| first-year comp. focal:biomass:adult comp. background | 0.00 | 0.00 | -0.01 | 0.01 |
| adult comp. focal:biomass:first-year comp. background | -0.01 | 0.01 | -0.02 | 0.01 |
| first-year comp. focal:biomass:first-year comp. background | 0.00 | 0.01 | -0.01 | 0.01 |
| fungicide:biomass:adult comp. background | 0.00 | 0.01 | -0.01 | 0.01 |
| fungicide:biomass:first-year comp. background | -0.01 | 0.01 | -0.02 | 0.01 |
| adult comp. focal:fungicide:biomass | 0.00 | 0.00 | 0.00 | 0.00 |
| first-year comp. focal:fungicide:biomass | 0.00 | 0.00 | 0.00 | 0.00 |
| adult comp. focal:fungicide:biomass:adult comp. background | 0.02 | 0.01 | 0.00 | 0.03 |
| first-year comp. focal:fungicide:biomass:adult comp. background | 0.00 | 0.01 | -0.01 | 0.01 |
| adult comp. focal:fungicide:biomass:first-year comp. background | 0.02 | 0.01 | 0.00 | 0.04 |
| first-year comp. focal:fungicide:biomass:first-year comp. background | 0.00 | 0.01 | -0.01 | 0.02 |
| random intercept: plot | 0.46 | 0.05 | 0.36 | 0.57 |
| sigma | 0.59 | 0.02 | 0.55 | 0.63 |

Notes: Prior distributions were: intercept – normal (location = 3, scale = 1), coefficients – normal (location = 0, scale = 1), random intercept – half Student's *t* (location = 0, df = 3, scale = 2.5)

**Table S24.** Estimated differences in disease severity (%) between untreated *M. vimineum* surrounding experimental plots and *M. vimineum* in plots where it was planted as the competitor (*N* = 192). Beta (log-link) generalized linear mixed-effects regression: disease severity ~ plant type * treatment * month, where plant type was either within plots or on the edge of plots. Positive values indicate severity on plants surrounding plots was higher than on plants within plots.

| Weeks post planting | Treatment | Estimate | Est. Error | l-95% CI | u-95% CI |
|---|---|---|---|---|---|
| 4 | control | -1.30 | 0.97 | -3.29 | 0.55 |
| 4 | fungicide | 0.30 | 0.90 | -1.45 | 2.08 |
| 8 | control | -0.04 | 0.94 | -1.87 | 1.84 |
| 8 | fungicide | 0.46 | 0.94 | -1.34 | 2.36 |
| 12 | control | -2.80 | 1.48 | -5.77 | 0.04 |
| 12 | fungicide | 0.63 | 1.17 | -1.62 | 3.00 |
| 16 | control | 14.47 | 3.92 | 6.73 | 22.03 |
| 16 | fungicide | 25.96 | 3.73 | 18.58 | 33.20 |

Notes: Prior distributions were: intercept and coefficients – normal (location = 0, scale = 1), phi – gamma (location = 0.01, scale = 0.01), random intercept – half Student's *t* (location = 0, df = 3, scale = 2.5)

**Table S25.** Timeline for experiment activities (month/day)

| Activity | 2018 | 2019 |
|---|---|---|
| *E. virginicus* seeds planted in flat | 2/26 and 3/2 | 3/25 |
| Transplanted *E. virginicus* to individual containers | 3/12–3/27 | 4/11–4/29 |
| *M. vimineum* seeds planted in flat | 3/26 and 3/30 | 3/28 |
| Transplanted *M. vimineum* to individual containers | 4/13–4/19 | 4/23–5/1 |
| Harvested *M. vimineum* litter within plots | N/A | 4/8–4/9 |
| Recorded overwinter survival of *E. virginicus* adults | N/A | 4/8–4/9 |
| Collected adult *E. virginicus* | 4/5 | 4/10, 5/8–5/10 |
| Plots raked and sprayed with herbicide | 4/27 | 4/10, 4/30 |
| Moved plants outside of greenhouse | 5/2 | 5/2 |
| Planted plots and watered | 5/9–5/11 | 5/8–5/10 |
| Collected plant growth data | 5/10–5/11, 5/15, 5/24, 6/5–6/7, 7/5–7/5 | N/A |
| Weed and water plots, replace dead plants | 5/15, 5/24, 7/7–7/9 (no replacements in July) | 5/15, 5/22, 5/27, 5/31, 6/6, 6/15, 6/23, 7/2, 7/3, 7/11, 7/23, 7/29–8/1, 8/16, 8/28–8/31, 9/12, 9/24–9/28 |
| Collected infection data | 6/5–6/7, 7/4–7/9, 8/1 - 8/2, 8/27–8/29, 9/24–9/26 | 5/7, 6/4–6/5, 7/2–7/5, 7/29–8/1, 8/28–8/31, 9/24–9/28 |
| Fixed planting position of #6 plots | N/A | 5/27 |
| Collected plot-level soil moisture data | 6/5–6/6, 10/22–10/23 | N/A |
| Collected site-level tree density and canopy data | 6/5–6/6 | N/A |
| Characterized site-level plant community | 6/6 | N/A |
| Sprayed with fungicide/water | 5/11, 6/7, 7/10, 8/3, 8/30, 9/27 | 5/10, 6/6, 7/5, 8/1, 8/31, 9/28 |
| Collected plot-level canopy cover data | 7/9 | |
| Placed data loggers | N/A | 7/2 |
| Collected *E. virginicus* seeds | 7/11, 8/1 - 8/2, 8/27–8/29, 9/24–9/26, 10/22 | 7/2–7/5, 7/29–8/1, 8/28–8/31, 9/24–9/28, 10/22–10/24 |
| Put seed collection bags on *M. vimineum* | 9/26–9/27 | 9/25–9/27 |
| Collected *M. vimineum* seed bags and biomass | 10/22–10/23 | 10/22–10/24 |
| Collected *E. virginicus* biomass | N/A | 10/22–10/24 |
| Collected data loggers | N/A | 10/22–10/24 |

**Table S26.** Parameter values for dynamical model

| Param. | Description | Fungicide | Control | Units | Source |
|---|---|---|---|---|---|
| $r_A$ | growth rate of annual | 0.039 | 0.037 | $day^{-1}$ | A |
| $r_F$ | fy perennial growth rate | 0.038 | 0.036 | $day^{-1}$ | A |
| $r_P$ | adult perennial growth rate | 0.029 | 0.031 | $day^{-1}$ | A |
| $\alpha_{AA}$ | annual–annual competition | 0.001 | 0.002 | $m^2\ g^{-1}$ | A |
| $\alpha_{AF}$ | fy perennial–annual competition | 0.000 | 0.000 | $m^2\ g^{-1}$ | A |
| $\alpha_{AP}$ | adult perennial–annual competition | $3.55e^{-4}$ | 0.000 | $m^2\ g^{-1}$ | A |
| $\alpha_{FA}$ | annual–fy perennial competition | 0.001 | $4.11e^{-4}$ | $m^2\ g^{-1}$ | A |
| $\alpha_{FF}$ | fy perennial–fy perennial competition | 0.000 | 0.000 | $m^2\ g^{-1}$ | A |
| $\alpha_{FP}$ | adult perennial–fy perennial competition | 0.001 | 0.008 | $m^2\ g^{-1}$ | A |
| $\alpha_{PA}$ | annual–adult perennial competition | 0.000 | 0.001 | $m^2\ g^{-1}$ | A |
| $\alpha_{PF}$ | fy perennial–adult perennial competition | 0.000 | 0.000 | $m^2\ g^{-1}$ | A |
| $\alpha_{PP}$ | adult perennial–adult perennial competition | 0.001 | 0.008 | $m^2\ g^{-1}$ | A |
| $\beta_{AA}$ | annual–annual transmission | $2.28e^{-5}$ | $2.97e^{-5}$ | $m^2\ g^{-1}\ day^{-1}$ | A |
| $\beta_{AF}$ | fy perennial–annual transmission | $1.53e^{-5}$ | $1.34e^{-5}$ | $m^2\ g^{-1}\ day^{-1}$ | A |
| $\beta_{AP}$ | adult perennial–annual transmission | $6.78e^{-6}$ | $1.84e^{-5}$ | $m^2\ g^{-1}\ day^{-1}$ | A |
| $\beta_{FA}$ | annual–fy perennial transmission | $3.51e^{-5}$ | $2.76e^{-5}$ | $m^2\ g^{-1}\ day^{-1}$ | A |
| $\beta_{FF}$ | fy perennial–fy perennial transmission | $1.74e^{-5}$ | $6.35e^{-6}$ | $m^2\ g^{-1}\ day^{-1}$ | A |
| $\beta_{FP}$ | adult perennial–fy perennial transmission | $4.51e^{-5}$ | $2.04e^{-5}$ | $m^2\ g^{-1}\ day^{-1}$ | A |
| $\beta_{PA}$ | annual–adult perennial transmission | $7.61e^{-5}$ | $3.53e^{-5}$ | $m^2\ g^{-1}\ day^{-1}$ | A |
| $\beta_{PF}$ | fy perennial–adult perennial transmission | 0 | $8.64e^{-6}$ | $m^2\ g^{-1}\ day^{-1}$ | A |
| $\beta_{PP}$ | adult perennial–adult perennial transmission | $3.29e^{-6}$ | $1.17e^{-6}$ | $m^2\ g^{-1}\ day^{-1}$ | A |
| $g_A$ | annual germination fraction | 0.779 | 0.761 | NA | A |
| $g_P$ | perennial germination fraction | 0.178 | 0.224 | NA | A |
| $c_A$ | seed conversion of annual | 69.0 | 76.9 | seed $g^{-1}$ | A |
| $c_F$ | fy perennial seed conversion | 7.78 | 8.47 | seed $g^{-1}$ | A |
| $c_P$ | adult perennial seed conversion | 20.7 | 19.0 | seed $g^{-1}$ | A |
| $e_A$ | max establishment fraction of annual | 0.999 | 0.989 | NA | A |
| $e_P$ | max establishment fraction of perennial | 0.994 | 0.944 | NA | A |
| $l_P$ | adult perennial survival fraction | 0.963 | 0.945 | NA | A |
| $n_P$ | max adult perennial biomass contribution | 0.418 | 0.410 | NA | A |
| $s_A$ | surviving seed fraction of annual | 0.150 | 0.000 | NA | B/A |
| $s_P$ | surviving seed fraction of perennial | 0.050 | NA | NA | C |
| $b_A$ | initial biomass of annual | 0.031 | NA | g | A |
| $b_F$ | fy perennial initial biomass | 0.004 | NA | g | A |
| $\gamma_A$ | sensitivity to litter of annual | 0.001 | NA | $m^2\ g^{-1}$ | D |
| $\gamma_P$ | sensitivity to litter of perennial | 0.001 | NA | $m^2\ g^{-1}$ | D |
| $h$ | conidia addition to litter | 5500 | NA | $m^2\ g^{-1}$ | D |
| $d$ | litter decomposition fraction | 0.590 | NA | NA | E |
| $a$ | conidia loss from litter | 0.500 | NA | $day^{-1}$ | O |
| $m_i$ | biomass mortality of any plant group | 0.002 | NA | $day^{-1}$ | A |
| $\beta_{iC}$ | litter transmission to any plant group | $1e^{-7}$ | NA | $m^2\ g^{-1}\ day^{-1}$ | O |
| $v_i$ | biomass virulence of any plant group | 0.001 | NA | $day^{-1}$ | O |
| $T$ | final day of growing season | 160 | NA | NA | A |

Notes: A = the experiment described in this paper, B = Redwood et al. 2018, C = Garrison and Stier 2010 (17), D = Benitez et al. 2022 (12), E = DeMeester and Richter 2010 (13), fy = first-year, O = expert opinion, $ae^{-b} = a \times 10^{-b}$
